# Dissecting Factors Underlying Phylogenetic Uncertainty Using Machine Learning Models

**DOI:** 10.1101/2023.09.20.558635

**Authors:** Ulises Rosas-Puchuri, Emanuell Duarte-Ribeiro, Sina Khanmohammadi, Dahiana Arcila, Guillermo Ortí, Ricardo Betancur-R

## Abstract

Phylogenetic inference can be influenced by both underlying biological processes and methodological factors. While biological processes can be modeled, these models frequently make the assumption that methodological factors do not significantly influence the outcome of phylogenomic analyses. Depending on their severity, methodological factors can introduce inconsistency and uncertainty into the inference process. Although search protocols have been proposed to mitigate these issues, many solutions tend to treat factors independently or assume a linear relationship among them. In this study, we capitalize on the increasing size of phylogenetic datasets, using them to train machine learning models. This approach transcends the linearity assumption, accommodating complex non-linear relationships among features. We examined two phylogenomic datasets for teleost fishes: a newly generated dataset for protacanthopterygians (salmonids, galaxiids, marine smelts, and allies), and a reanalysis of a dataset for carangarians (flatfishes and allies). Upon testing five supervised machine learning models, we found that all outperformed the linear model (p < 0.05), with the deep neural network showing the best fit for both empirical datasets tested. Feature importance analyses indicated that influential factors were specific to individual datasets. The insights obtained have the potential to significantly enhance decision-making in phylogenetic analyses, assisting, for example, in the choice of suitable DNA sequence models and data transformation methods. This study can serve as a baseline for future endeavors aiming to capture non-linear interactions of features in phylogenomic datasets using machine learning and complement existing tools for phylogenetic analyses.

## 1. Introduction

The inference of phylogenetic trees aids in our understanding of the evolutionary history of species, and it has found applications in research areas such as ecology, effects of climate change analyses, and protein structure prediction (Felsenstein 2004; Isaac et al. 2007; Kratzer et al. 2014; Li and Weins 2021; Jumper et al. 2021). The advent of genomic datasets has further improved our capacity to consistently elucidate sections of the tree of life previously unresolved. However, due to underlying biological processes and methodological or analytical factors, phylogenetic inference can yield incongruent or inconsistent tree topologies even using genomic-scale datasets. Biological processes that produce misleading phylogenetic results —if not accounted for— include incomplete lineage sorting, introgression, paralogy and natural selection (Maddison 1997; De Queiroz 2007; Kapli et al. 2020). While it is presently possible to model such biological processes (Simão et al. 2015; Solís-Lemus and Ané 2016; Zhang et al. 2018), they often assume that no analytical factors that may cause estimation error are at play. However, in reality, the results of phylogenomic analyses stem from a combination of uncertainty originating from both biological processes and analytical factors (Cai et al. 2020; Steenwyk et al. 2023).

Analytical factors including properties or features of the data vary and impact phylogenetic reconstructions in distinct ways (Kumar et al. 2012; Betancur-R. et al. 2013; Susko and Roger 2021; Kapli, Yang, and Telford 2020). Missing taxa provides a relevant example, as studies have reported that increasing taxon sampling can enhance inference stability (Heath, Hedtke & Hillis 2008; Betancur-R. et al. 2019; Singhal et al. 2021). But there are also cases where specific taxa consistently destabilize the inference (Aberer, Krompas and Stamatakis 2013). Additionally, datasets with an insufficient number of sites can produce low-quality gene trees, which can introduce subsequent errors when inferring the species tree from them (Springer and Gatesy 2016; Molloy and Warnow 2018). Model misspecification, another example, can introduce bias when dealing with a large number of sites, as is common with current genomic datasets (Kumar et al. 2012; Doyle et al. 2015). While the above-mentioned sources of incongruence can be addressed individually, proposed corrections often require computationally demanding approaches or more sampling efforts (Xi, Liu, and Davis 2016; Kapli et al. 2020; Steenwyk et al. 2023). For example, it might be necessary to increase the sequencing budget to obtain more markers, increase the sampling effort to collect missing species, or employ more thorough runs for model-selection software. Additionally, analytical factors may covary or interact with one another (Xing Xing Shen, Salichos, and Rokas 2016; Mongiardino 2021), which adds complexity to the decision-making process about which factors to account for. Therefore, understanding these factors beforehand can enhance the efficiency of phylogenetic inference. This is achieved by either avoiding computationally intensive runs — thereby aligning with green computing principles (Kumar 2022) — or by applying necessary corrections.

To understand how these factors affect phylogenomic datasets, one approach involves close examination of values from a predefined set of features relevant to phylogenetic inference within the dataset. The selection of these features is based on their correspondence with analytical factors, as suggested by previous studies. An alternative approach leverages the associations of these features with specific support metrics, such as likelihood scores, bootstrap support, or p-values from topology tests, for specific phylogenetic hypotheses (Xing-Xing Shen, Steenwyk, and Rokas 2021; Steenwyk et al. 2023). Based on these associations, researchers can dissect the relative importance of these factors in causing phylogenetic incongruence (Xing-Xing Shen, Steenwyk, and Rokas 2021) and then formulate future corrections to minimize bias and error. However, studies employing above outlined approaches often evaluate the features independently (e.g., Xing-Xing Shen, Steenwyk, and Rokas 2021; Duchêne et al. 2021; Gosselin et al. 2021; Cunha, Reimer, and Giribet 2022) or assume that there exists an underlying linear relationship among them (Xing Xing Shen, Salichos, and Rokas 2016; Vankan, Ho, and Duchêne 2021). Although linearity might serve as a suitable initial assumption in the modeling of biological relationships, it is often not a biologically realistic one (Strogatz 2018). For instance, various systems — such as predator-prey dynamics (Lotka 1925; Volterra 1927), biochemical processes (Sel’Kov 1968), the spread of infectious diseases (Kermack and McKendrick 1991), and in certain cases, phylogenetic regressions (Quader et al. 2004) — exhibit highly non-linear feature interactions. Then, in phylogenomic datasets, a more efficient approach could involve implementing a methodology that relaxes the assumption of linearity while simultaneously considering feature interactions (Ronquist and Huelsenbeck 2003; Steel 2005; Zou et al. 2020). Unfortunately, as of now, such an option does not exist due to challenges of constructing and parameterizing these models from first principles, as we usually lack full knowledge about the theoretical distribution of these features and how they interact with each other (Susko and Roger 2021). Hence, there is a need to develop non-linear models that do not rely on prior assumptions about the distribution and interactions of the features.

With the emergence of large phylogenomic datasets containing thousands of markers (Hughes et al. 2018; Álvarez-Carretero et al. 2022; Harvey et al. 2020), we now have the opportunity to extract a substantial number of feature instances from each marker, including alignment information (e.g., GC-content, number of taxa) and tree information (e.g., average bootstrap support, sum of branch lengths). This presents an unprecedented opportunity to train machine learning models that can potentially learn complex non-linear relationships within the phylogenomic dataset without imposing a fixed set of probability distribution families to construct the model (i.e., non-parametric models) (Hastie et al. 2009; Murphy 2012; Bokulich et al. 2018). Machine learning models, capable of identifying complex non-linear relationships, have recently found applications in various phylogenetic analyses, including fast phylogenetic inference (Azouri et al. 2021, 2023), phylogenetic placement (Jiang, Tabaghi, and Mirarab 2022), learning phylodynamic parameters (Voznica 2021), and selecting biogeographical models from simulated data (Smith et al. 2017; Burbrink and Gehara 2018). While most of these studies focus on prediction tasks, exploring the interactions of features in specific phylogenetic datasets leveraging these non-linear models remains largely unexplored.

Here, we propose the use of machine learning models to correlate the set of features known as factors that introduce biases and errors in phylogenomic inference with the phylogenetic signal within a phylogenomic dataset. This will allow the features to interact non-linearly, yielding valuable insights from the derived machine learning models. Because phylogenomic datasets contain data from hundreds or thousands of genes, we can extract numerous feature instances where machine learning models can function effectively. There are multiple ways to obtain the relative support a given phylogenetic hypothesis receives from each of the genes. For example, we can derive the signal based on likelihood scores (Xing Xing Shen, Hittinger, and Rokas 2017), concordance factors (Minh, Hahn, and Lanfear 2020), or quartet scores (Zhang et al. 2018; Xing-Xing Shen, Steenwyk, and Rokas 2021). Another alternative, which we implement here, is the use of Gene-Genealogy Interrogation (GGI) (Arcila et al. 2017) to assess the relative support that alternative hypotheses receive based on p-values from the distribution of site log-likelihoods (Shimodaira 2002), as a label to gene marker features. Although machine learning models can be complex and are often seen as “black boxes,” we can improve their “explainability” by using feature importance analyses (Breiman 2001; Lundberg and Lee 2017), which assess how each feature influences the overall model by permuting the feature structure.

This study utilizes supervised learning models (Hastie et al. 2009) that, although non-parametric, differ in their optimization strategies to minimize the error between the predicted and observed labels. Examples include Support Vector Machines (Cortes and Vapnik 1995), which fit a linear function after transforming the feature space into a potentially infinite-dimensional one. Random Forest and Gradient Boosting (Breiman 2001; Chen and Guestrin 2016) use multiple decision trees (Breiman et al. 1984) for making predictions. Deep Neural Networks (LeCun, Bengio, and Hinton 2015) and their architectural variants, such as Autoencoders, deploy networks whose nodes are organized into layers and whose edges are weighted. This allows them to extract increasingly abstract data representations as the number of hidden layers increases (Goodfellow, Bengio, and Courville 2016). The versatility of these models allows for the adjustment of multiple hyperparameters, resulting in the formation of varied architectures suited to the specific dataset. Notably, the requirement for hyperparameter adjustment tends to decrease as the dataset size increases (Ng 2004; Goodfellow, Bengio, and Courville 2016).

Here, we applied the outlined machine learning models to two phylogenomic datasets collected to resolve controversial relationships among fish clades. Rather than simulating the phylogenetic features, we refrained from using such an approach due to the typical uncertainty regarding the accurate distribution and interplay of these features. The first dataset encompasses “protacanthopterygian” fishes, which was newly generated for this study. Protacanthopterygii is a higher-level, early-branching euteleost lineage that was proposed 57 years ago (Greenwood et al. 1966) and has undergone at least 13 changes in its circumscription (Ishiguro, Miya, and Nishida 2003; R. R. Betancur et al. 2017), but previous analyses have not adequately dealt with either comprehensive species coverage or genome-wide genetic sampling. The second dataset was recently generated for fishes in the series Carangaria (i.e., flatfishes, billfishes, remoras, and allies; Duarte-Ribeiro et al., under review), which we re-analyzed here. The most contentious relationship in Carangaria concerns the monophyly of flatfishes and the evolutionary implication of single versus dual evolutionary origins of their asymmetric body plan (Betancur-R. et al., 2013; Harrington et al., 2013; Campbell et al., 2013; Lü et al., 2021). We constructed a matrix with 40 relevant features for phylogenetic inference based on both alignment and tree information. We fitted five supervised machine learning models, obtained the best-fitted model by using cross-validation, and assessed feature importance for each of the datasets analyzed. Ultimately, our goal is to use machine learning to dissect factors underlying phylogenetic uncertainty by accounting for the non-linearity of phylogenomic features.

## 2. Materials and Methods

### 2.1. Case study datasets

We generated a new dataset for Protacanthopterygii (77 species and 1023 exons) and reanalyzed a recently generated dataset for Carangaria (395 species and 991 exons; Duarte-Ribeiro et al., under review), both of which are part of a large initiative—the FishLife project—aimed at estimating phylogenomic trees for over six thousand fish species based on exon capture approaches. Ever since the pre-cladistic definition of Protacanthopterygii by Greenwood et al. (1966), this clade was conceived as encompassing various taxa deemed as “precursors” of acanthopterygians. Although this concept is no longer supported, numerous alterations to its membership and interrelationships have been suggested over the past 57 years, totaling at least 13 different proposed changes. A major problem with previous studies is that they have not adequately dealt with both comprehensive species coverage and genome-wide genetic sampling. We chose the Protacanthopterygii dataset with the goal of analyzing phylogenomic evidence to assess support for alternative hypotheses regarding the relationships within this group and among its close allies. Our objective is also to pinpoint variables that could provide explanations for, or enhance the resolution of, the persistently debated early-branching euteleost lineages (e.g., Hughes et al. 2018). We also obtained publicly available sequences to expand the taxonomic coverage for the Protacanthopterygii dataset to 123 species (see Table S1). We chose the Carangaria dataset for reanalysis here given that previous studies have indicated that base-composition heterogeneity is a major factor affecting exonic datasets generated for this clade (Betancur-R. et al., 2013; Ribeiro et al., 2018), with resulting phylogenetic analyses often (though not always) failing to resolve the monophyly of flatfishes, one of the largest clades in Carangaria. Such conflicting results have important evolutionary implications as they may either imply that the flatfish asymmetric body plan has a single (see Betancur-R. et al., 2013; Harrington et al., 2013) or dual (see Campbell et al., 2013; Lü et al., 2021) origin. See Tables S1 and S2 for the complete list of species for both datasets analyzed.

### 2.2 Exon capture, assembly, and quality control

We extracted high-yield DNA for 62 species for the Protacanthopterygii dataset (orders Salmoniformes, Esociformes, Argentiniformes, Osmeriformes, Stomiatiformes, Galaxiiformes), also including a subset of 15 outgroup species (orders Lepidogalaxiiformes, Alepocephaliformes, Clupeiformes, and Neoteleostei) and generated enriched genomic libraries using the “backbone 1” and “backbone 2” probe sets of Hughes et al. (2021). Arbor Biosciences (Ann Arbor, MI) performed library preparation using dual-round capture protocol (Li et al. 2013), and the University of Chicago Genomics Facility sequenced the resulting libraries using a HiSeq4000 to obtain pair-end 150 bp reads, with 190 samples multiplexed per lane. We filtered out raw reads containing adaptor contamination (with >15 bp matched to the adaptor sequence), >10% of low-quality bases (quality score <10), or > 2% unknown bases. We then added publicly available sequences, and we assembled all filtered reads using an optimized version of the FishLifeExonCapture pipeline (github.com/ulises-rosas/FishLifeExonCapture; Hughes et al., 2021). Briefly, this pipeline uses Trimmomatic v0.36 (Bolger et al., 2014) to trim raw FASTQ files based on the read quality function, BWA-MEM (Li and Durbin, 2009) to map trimmed reads against reference sequences, SAMtools (Li et al., 2009) to remove PCR duplicates, and aTRAM (Allen et al., 2017) to extend mapped reads after an initial assembling step with Velvet (Zerbino and Birney, 2008) and a final stage with Trinity (Haas et al., 2013). CD-HIT (Fu et al., 2012) is then used to identify identical sequences and exonerate (Slater and Birney, 2005) to define open reading frames upon collecting sequences for all target exons. Finally, the pipeline uses MACSE (Ranwez et al., 2011) to merge sequences for each exon for all taxa and to align them in by taking into account the underlying codon structure of the protein-coding sequences. The final Protacanthopterygii dataset assembled includes 1133 exon alignments for 123 species (see Results). We also assembled the Carangaria dataset from the previous study in a similar manner. This dataset was generated using the “Carangaria” probe sets of Hughes et al. (2021) and consisted of 991 exon sequences for 395 species, including 12 non-carangarian outgroups. After assembly, we applied a quality control pipeline to the datasets to address potential sources of contamination, mislabeling, sequence error, or other confounding features of the data. We based this pipeline on the conceptual framework proposed by Simion et al. (2017) and Arcila et al. (2021), by accounting for missing data, taxon mislabeling, and taxon misplacement. We integrated the steps mentioned above into a Python-based command-line interface called “fishlifeqc” (available at: github.com/ulises-rosas/fishlifeqc); see Supplemental Information S1 for more details.

### 2.3 Phylogenomic inference

All downstream analyses applied to both datasets are based on trees estimated using both concatenation and multispecies coalescent approaches. However, for this study we only inferred phylogenetic trees for the Protacanthopterygii dataset, and instead reused trees for the Carangaria dataset from Duarte-Ribeiro et al. (under review) study, which were estimated following similar procedures described below.

We used IQ-TREE version 2.0.6 (Nguyen et al., 2015) to estimate gene trees and concatenation-based phylogenies under the maximum likelihood (concatenation-based ML hereafter) approach. We first generated concatenated gene alignments using “concatenate.py” (a utility of “fishlifeqc”) for Protacanthopterygii, and then used ModelPartition (Kalyaanamoorthy et al., 2017) to select the best-fit partitioning scheme (using the “-p” option) based on by-gene and by-codon *a priori* partitions. This step generated 258 partitions, including best-fit evolutionary models for each partition. Concatenation-based analyses used the following IQ-TREE options: `“iqtree2 -s [alignment file] -p [partition file] -T [Number of cores] --ufboot 1000”`. We then used Ultrafast Bootstrap (Hoang et al., 2018) to assess edge support (set to 1000 using the “--ufboot 1000” option).

We also used IQ-TREE based gene trees as input for multispecies coalescent (MSC hereafter) analyses as implemented in ASTRAL-III (Zhang et al., 2018). To minimize artifacts caused by gene tree estimation error, we collapsed edges with low support (aLRT = 0, Anisimova and Gascuel, 2006), following recommendations by Simmons and Gatesy (2021). By default setting, ASTRAL-III automatically used the quartet score, which is the proportion of quartet trees induced from input gene trees supporting the species tree, to assess the edge support. Each dataset produced two major alternative hypotheses, one under the ML approach and the other under the multispecies coalescent approach.

These competing hypotheses differed in key aspects of the evolutionary history of each clade (see details under Results, Fig 1, and Fig. S1). For instance, the Protacanthopterygii trees differed in the relative placement of the order Argentiniformes, and the Carangaria dataset on the monophyletic status of flatfishes (comprised by suborders Psettodoidei and Pleuronectoidei; Fig. 1). Subsequent analyses assess the relative support that each competing hypotheses receive from individual exon alignments.

**Figure 1.**
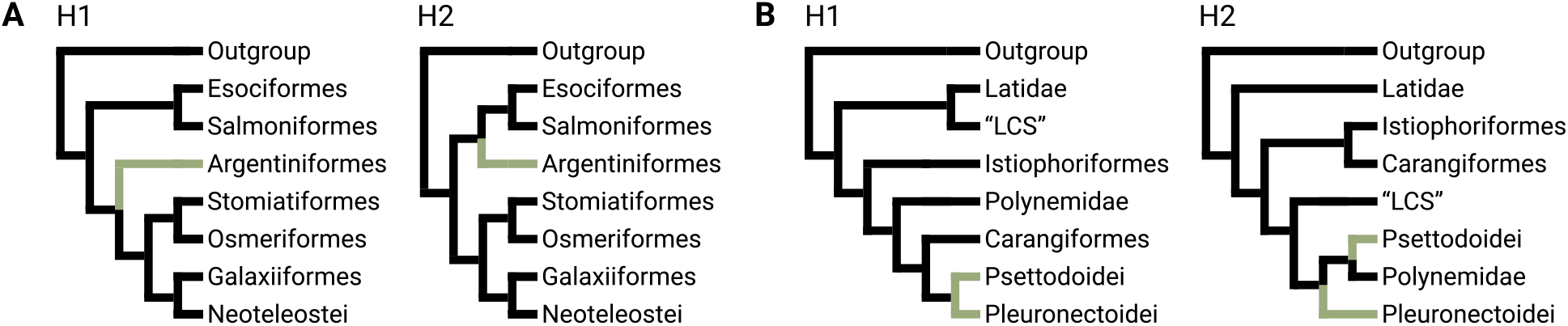
Phylogenetic trees obtained for Protacantopterygii (**A**) and Carangaria (**B**). In **A**), H1 is the concatenation-based ML tree and H2 the MSC tree (also see Results and Fig. 3). These trees differ in the placement of Argentiniformes (highlighted with green). In **B**), H1 is the MSC tree and H2 the concatenation-based ML tree (see Fig. S1). While multiple differences exist between these two alternative trees, a major discrepancy comprises the monophyletic status of flatfishes which includes the suborders Psettodoidei and Pleuronectoidei. In both hypotheses, these suborders are highlighted in green. The acronym “LCS” designates the clade consisting of Lactariidae, Centropomidae, and Sphyraenidae.

### 2.4 Gene-Genealogy Interrogation (GGI)

We applied GGI (Arcila et al., 2017) to quantify the amount of phylogenetic signal in the loci supporting each competing hypothesis. Briefly, GGI steps include defining competing hypotheses, building constraint trees by pruning the set to taxa to match those in individual alignments, estimating constrained ML gene trees under the GRT+GAMMA model in RAxML, estimating site log-likelihoods (SLL) for each constrained gene tree, and conducting AU-tests using the SLL as input to assess the relative support that each hypothesis receives from individual genes. Complex datasets (i.e., datasets that presumably violate model assumptions; Pupko & Mayrose 2020) can affect the convergence of the numerical optimization of the α parameter of the GTR+GAMMA model (Jermiin et al. 2017, Naser-Khdour et al. 2019). Thus, for each exon, instead of putting both constraints in a single file, which makes the initial α value for the second constraint to be the optimized by the α value from the first constraint (see the source code: https://github.com/stamatak/standard-RAxML/blob/master/axml.c#L10493-L10514, last accessed June 20, 2023), we obtained SLL for each constraint separately (i.e., one constraint tree per file). This final step was incorporated to tackle certain concerns raised by Simion et al. (2020) regarding potential biases stemming from RAxML in estimating SLL (site likelihoods) for GGI specifically, and for topology tests more generally. These steps were integrated into a Python-based command line interface called “ggpy” (available from github.com/ulises-rosas/ggpy). The output of each AU test from above provides a preferred gene tree and a rank of preferences for the alternative hypotheses for each exon, based on their likelihood score, each weighted by its p-value.

For the Protacanthopterygii dataset, we tested two hypotheses obtained from the previous step (concatenation-based ML and MSC trees) as well as 103 other possible rooted topologies (i.e., orders Lepidogalaxiiformes, Alepocephaliformes, and Clupeiformes composed the outgroup in the root), for a total of 105 trees based on all possible relationships among five (well-supported) subclades (see Table S3). These subclades are: (1) Esociformes + Salmoniformes (“Eso-Salmo” hereafter), (2) Osmeriformes + Stomiiatiformes (“Osme-Stomia” hereafter), (3) Neoteleostei, (4) Galaxiiformes, and (5) Argentiniformes. The GGI analyses (see Fig. S2) show that the top two trees favored are also the top trees resolved with concatenation-based ML and MSC tree inference (see previous section). We treated these top two hypotheses, derived from phylogenetic inferences, as the main hypotheses. However, only 114 and 55 loci achieved rank 1 (i.e., the lowest AU p-value) for these hypotheses, and neither was statistically significant (i.e., AU p-value > 0.05). To enhance the statistical power of the AU p-value associated with the concatenation-based ML and MSC hypotheses, we conducted the GGI focusing solely on these two topologies, mirroring the approach of Shen, Hittinger & Rokas et al. (2017). The Carangaria dataset includes all 30 families comprised in this evolutionary radiation (Ribeiro et al., 2018; Duarte-Ribeiro et al., under review), yielding a total of ~4.95e^38^ possible interfamilial topologies. Because running GGI with ~4.95e^38^ topologies is computationally intractable, we instead used the top two hypotheses obtained with phylogenetic inference. For both datasets, the best tree favored among the two competing hypotheses tested for each gene alignment was used as input for the machine learning model component (next section). After the above GGI runs for each dataset, each exon has two AU-based p-values associated with a given hypothesis. We used these p-values as label input for the machine learning models.

### 2.5 Machine learning models

To explore the relationship between phylogenetic signal and exon alignment properties associated with a given tree, we applied five supervised machine learning models (Hastie et al. 2009) for multi-output regression analyses where the number of features includes 40 relevant properties for phylogenetic inference (Table S4), each with 2 alternative labels for each exon alignment. In the case of labels, more generally, if we let an exon alignment *i* have two AU p-values *p*_*i*,1_ and *p*_*i*,2_ for the first and second hypothesis, respectively, then the label for exon alignment *i* will be the row vector [*p*_*i*1_ *p*_*i*,2_]. For our feature set, we derived them from gene alignments and gene trees, incorporating 40 previously proposed properties deemed relevant for phylogenetic inference as per Shen, Salichos, & Rokas (2016) and Haag et al. (2022). Several studies have identified these features as being connected to factors that introduce uncertainty in phylogenetic inference. For instance, a high variance in GC content can be linked to violations of DNA sequence model assumptions (Kolaczkowski and Thornton 2004; Kapli et al., 2020), while the number of variable sites correlates with the desired noise-to-signal ratio (Zhang et al. 2021; Di Franco et al. 2022). Although there is no established consensus on the best set of features for machine learning models in phylogenetic analyses, supervised machine learning models employ regularization techniques during training to adjust feature importance and mitigate overfitting. These techniques include the tree depth in decision tree-based models and the dropout rate of neurons in Deep Neural Networks. Therefore, a comprehensive feature set is beneficial as the supervised machine learning model should be able to discern the most relevant patterns in the dataset. These can subsequently be identified through feature importance analyses, as discussed later.

Each dataset has a matrix for features and labels up to this point. We then split the above matrices into training and testing sets, consisting of around 66% and 33% of matrices rows, respectively (Fig. 2). We used both sets to perform hyperparameter tuning over the machine learning models. Since we have two labels per row, all machine learning models we tested were multioutput regressions, whose objective functions involve error minimization (i.e., Mean Squared Error, MSE) in the following form:

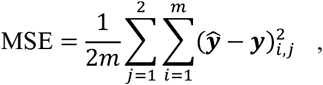

where 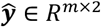 and ***y*** ∈ ***R***^*m*×2^ are the predicted and observed values, respectively, and *m* is the number of rows of a given dataset.

**Figure 2.**
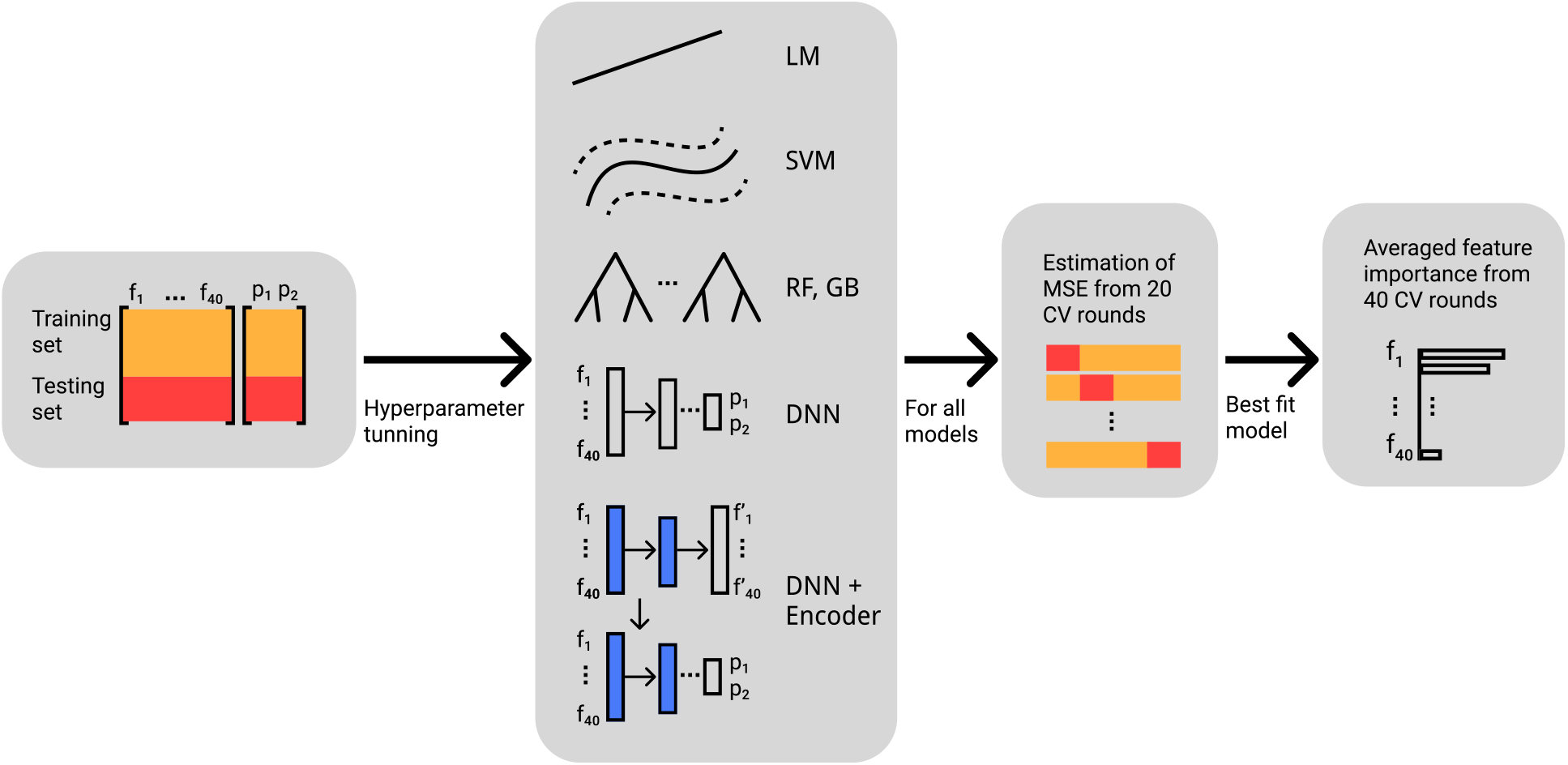
The pipeline starts with a case study dataset (e.g., Protacanthopterygii, Carangaria). i) Each dataset is first split into training (~66% of rows) and testing (~33% of rows) sets. ii) Then all considered models go through a hyperparameter tuning process. iii) We define the best-fit model for a given dataset as the model with the lowest average MSE of 20 cross-validations (CV) rounds. iv) With the best-fit model, we obtain the average feature importance of 40 CV rounds. LM: Linear Model; SVM: Support Vector Machine; RF: Random Forest; GB: Gradient Boosting; DNN: Deep Neural Network; and DNN + Encoder: Deep Neural Network with Encoder layers.

While all these models are non-parametric, they differ in the optimization strategy, which is how the model minimizes the error function. Support Vector Machines (SVM hereafter) (Cortes and Vapnik 1995), for example, assume a single optimal point (i.e., convex optimization) when the feature space is extended to a high-dimensional space. The advantage of this model is that it has at most two hyperparameters that need to be tuned. On the other hand, decision tree-based and neural network-based methods can handle local optimality (i.e., nonconvex optimization). Examples of decision tree-based models are Random Forest (RF; Breiman 2001) and Gradient Boosting (GB; Chen and Guestrin 2016). RF takes random sampling with replacement of the data to construct an ensemble of decision trees (i.e., unconnected acyclic graphs) (Breiman et al. 1984) and then averages the predictions from the ensemble of decision trees to make a prediction. GB sequentially combines several weak learners (i.e., decision trees with accuracy just better than random), while also accounting for predecessor residual errors, to form a strong learner (Hastie et al. 2009).

There exist many types of neural network-based models, including those used for text classification and image recognition. Most of them solely depend on Deep Neural Networks (DNN hereafter; LeCun, Bengio, and Hinton 2015; Goodfellow, Bengio, and Courville 2016). The simplest DNN architecture can be seen as a connected cyclic graph with three subsets of nodes: an input layer, a hidden layer, and an output layer. The edges, which are weighted, only exist between subsets but not within subsets. The weights from all edges are randomly initialized, and each node in the input layer represents a feature from instances. The value from each node that is not in the input layer is obtained from the application of a non-linear activation function (e.g., ReLU function) (Agarap 2018) over the weighted sum (plus a bias term) from the previous layer (Géron 2019). Backpropagation iterations (LeCun et al. 1989) update the weights from all edges based on the minimization of an error function between the value of the output layer (i.e., prediction) and the observed label. It has been pointed out that a simple DNN is equivalent to SVM with the number of nodes in the hidden layer approaching infinity (Jacot, Gabriel, and Hongler 2018; Domingos 2020). We briefly described the simplest DNN architecture, but there are many hyperparameters to tune (e.g., the number of nodes in the hidden layer, the number of hidden layers, etc.), allowing different architectures (e.g., Szegedy et al. 2015; He et al. 2016). One variation of the DNN architecture is Variational Autoencoders (Kingma and Welling 2013), which learn low dimensional embedding of noisy data. Its architecture is symmetrical with respect to the central layer, resembling the shape of a sand clock. It employs unsupervised training to learn the features itself. The encoder, comprising the first half of layers, converts the inputs into a latent representation at the central layer. The decoder, composed of the second half of layers, reconstructs the internal representation to generate outputs while considering some Gaussian noise. Since the first half layers learn low-level structures from the input, they can be transferred to create a new deep neural network for the actual task, a process known as transfer learning (Géron 2019).

We did not perform hyperparameter tuning for the Linear Model (LM) because it has an analytical solution through ordinary least squares. We used the Scikit-Learn python package (Pedregosa et al. 2011) to get the LM model, SVM, RF, and GB machine learning models, and its “RandomizedSearchCV” function to obtain the best set of hyperparameters given the dataset. We set two rounds of cross-validations per set of hyperparameters tested.

Most of the hyperparameters tuned directly affected the model overfitting prevention. For the SVM model, we tuned the C hyperparameter (i.e., 1 – 30, under a reciprocally continuous random variable) that controls the width of the margin error, and we also tuned the gamma hyperparameter (i.e., scale 1, under an exponentially continuous random variable) from the Radial Basis Function Kernel, which controls the relative importance of higher order terms in the expansion of the exponential function. For the RF model, we tuned the maximum depth of decision trees (i.e., 5 – 100), the minimum required sample size to split a node into leaf nodes (i.e., 2 – 50), and the maximum number of leaf nodes (i.e., 100 – 250). For the GB model, we used the efficient implementation of the model from the XGBoost package (Chen and Guestrin, 2016), and then tuned the maximum depth of decision trees (i.e., 3 – 15), the proportion of subsampled features (i.e., 0.1 – 1), gamma values (i.e., 1 – 10), the minimum leaf weight (i.e., 0.1 – 10), and the learning rate (i.e., 1e^-7^ – 9e^-1^).

We implemented DNN architectures and DNN including layers from a Variational Autoencoder architecture (DNN + Encoder hereafter) using TensorFlow through the high-level deep learning Keras API. We tuned their hyperparameters by using the “BayesianOptimization” function from the KerasTuner python package (O’Malley et al., 2019). The tuned hyperparameters were the number of hidden layers (i.e., 4 – 10), the number of neurons per layer (i.e., 5 – 100), the proportion of dropout neurons per layer (i.e., 1e^-4^ – 0.9), the learning rate (i.e., 1e^-4^ – 1e^-1^), and the decay rate of the learning rate (i.e., 1e^-7^ – 9e^-1^). As fixed hyperparameters, we used the SELU activation function (Klambauer et al. 2017), initialized network weights with the LeCun strategy (Glorot & Bengio 2010), and used the Nesterov Accelerated Gradient as an optimizer (Nesterov 1987). The Supplemental Material contains the complete list of selected hyperparameters of the above models for each dataset.

In the case of DNN + Encoder, we pursued a transfer learning approach. Briefly, we took out the Encoder layers from a Variational Autoencoder architecture and transferred them as base layers into a new DNN architecture (Fig. 2). We ran the Variational Autoencoder with 100,000 iterations, and the architecture involved a descending number of units from the input layer (40 units) to the central layer (20 units), and an ascending number from the central layer to the output layer (40 units). The difference in the number of units between hidden layers was 5. The gaussian sampling size for the central layer was 10, and its parameters (i.e., mean and variance) were updated for each iteration. We used the root mean squared propagation (RMSProp, Hinton, Srivastava & Swersky 2012) as the optimizer. The MSE for the Autoencoder architecture reached values below 0.16. Once we transferred the Encoder layer into a new DNN architecture (i.e., DNN + Encoder), we did not update their weights anymore, and hyperparameter tunning followed as above for only the new hidden layers.

We selected the best machine learning model for each dataset by assessing the average MSE after 20 rounds of cross-validation over different sample subsets (Fig. 2). For each subset, the proportion of training and testing samples was approximately 66% and 33%, respectively.

### 2.6 Feature importance analysis

Once we obtained the best-fit machine learning model for each dataset, we characterized the feature importance profile within each. If we let MSE_*i*_ be the MSE of the best-fit model at cross-validation round *i*, and 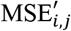 be the MSE of the best-fit model after randomly permuting values of feature *j* at cross-validation round *i*, then we can define the feature importance FI_*i,j*_ of feature *j* at cross-validation *i* as:

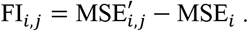

The above random permutation approach was first proposed for Random Forest models by Breiman (2001) and then Fisher, Rudi and Dominici (2018) extended it into a model-agnostic version. We used the eli5 python package to obtain FI_*i,j*_ values. We set the number of permutations to 100 to approach the 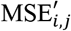 value. Thus, we can define the feature importance FI_*j*_ of feature *j* ∈ {1, …, 40} as the following average over cross-validation rounds:

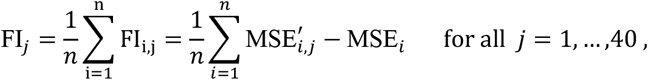

where *n* is the number of cross-validation rounds (*n* = 40 in our case). We increased the number of cross-validations rounds to account for the random effects of permutation. We can define the feature importance profile in a given dataset as the descending order of the set {FI_*j*_ | 1 ≤ *j* ≤ 40}.

## 3. Results

### 3.1 Dataset properties, Phylogenomic analyses, and GGI

We assembled 1133 exon alignments for the Protacanthopterygii dataset for 123 species (99 ingroup and 24 outgroup species). Raw reads generated in this study have been deposited in GenBank, under the following Bioproject Accession ID: PRJNA1013549. After applying a number of quality control steps, we retained 1023 (319,386 aligned sites) and 991 (457,167 aligned sites) exon alignments for the Protacanthopterygii and Carangaria (see details in Duarte-Ribeiro et al., under review) datasets, respectively. The length of exon alignments obtained is as follows: Protacanthopterygii dataset, 93–2349 sites (mean = 367.4, std = 276.1); Carangaria dataset, 117–5049 sites (mean = 461.3, std = 387.5).

Fig. 3 and Fig. S1 show both concatenation-based ML and MSC trees for the Protacanthopterygii and Carangaria datasets, respectively. For Protacanthopterygii, Argentiniformes was the sole major clade with a variable position among the top two competing trees (H1 and H2). This lineage grouped with Esociformes + Salmoniformes in the MSC tree (H2), but was resolved closer to Osmeriformes, Stomitatiformes, Galaxiiformes, and Neoteleostei in the concatenation-based ML tree (H1). For Carangaria, the ML and MSC trees include many more incongruent clades, but a remarkable difference relates to the monophyly of Pleuronectiformes. This clade is resolved as monophyletic in the MSC tree (H1) but is deemed paraphyletic in the concatenation-based ML tree (H2), with Psettodoidei being closer to the (non-flatfish) Polynemidae than to pleuronectoid flatfishes (see Fig S1; Duarte-Ribeiro et al., under review).

**Figure 3.**
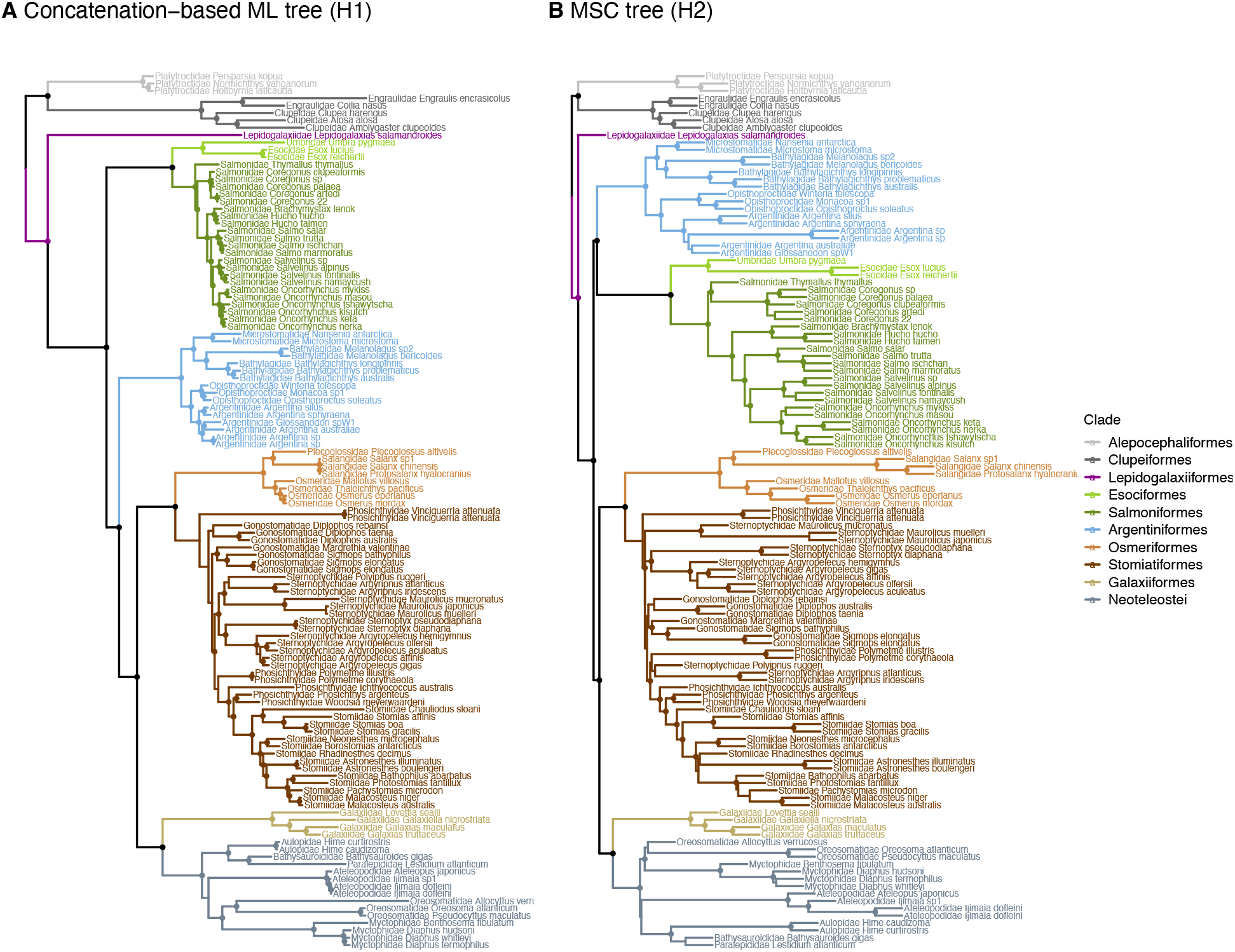
Two major phylogenetic trees estimated with the Protacanthopterygii dataset. **A)** concatenation-based ML tree obtained with IQ-TREE (H1); **B)** MSC tree obtained with ASTRAL (H2). Note the variable position of Argentiniformes (in sky-blue). Nodes with filled circles indicate ultrafast bootstrap value ≥ 99 % (H1) or quartet score of 1 (H2).

Neither dataset shows a noticeable separation between H1 and H2 in terms of AU p-values (see Fig. S3), although H1 is the most frequently favored tree topology for Protacanthopterygii (554 exons vs. 463 exons for H2; we discarded six exons due to long-running times of GGI) while the same is true for H2 in Carangaria (582 exons vs. 409 exons for H1). Average AU p-values for H1 and H2 were 0.48 and 0.46, respectively, for the Protacanthopterygii dataset, and 0.44 and 0.55, respectively, for the Carangaria dataset (see Fig. S4). For the Protacanthopterygii dataset, the average of the rank 1 AU p-value (i.e., lowest AU p-value) among exon alignments was 0.27, of which only 98 had an AU p-value < 0.05. Likewise, in the Carangaria dataset, the average of the rank 1 AU p-value among exons was 0.22, of which 172 had an AU p-value < 0.05. For both datasets, no evidence of noticeable separation of H1 and H2 in terms of AU p-values was found between exons with a large number of sites (see Fig. S3).

### 3.2 Machine Learning and Feature Importance

The best-fit model was DNN for both datasets according to the average MSE (Fig. 4). Conversely, the model with the highest MSE average was LM for both datasets. For both datasets, the difference in MSE between the DNN and LM models was statistically significant according to the Wilcox Rank Sum Test (Fig. 4). Furthermore, LM was the only model whose MSE had a significant difference relative to the rest of the models (p-value= 0.0304 for Protacanthopterygii; p-value= 0.0203 for Carangaria). A common attribute of all non-LM models is that they allow non-linear relationships among tested features. While the DNN has less bias compared to LM for both datasets, the MSE variance of the DNN for the Protacanthopterygii dataset (3.07e^-5^) was higher than that for the Carangaria dataset (2.26e^-5^).

**Figure 4.**
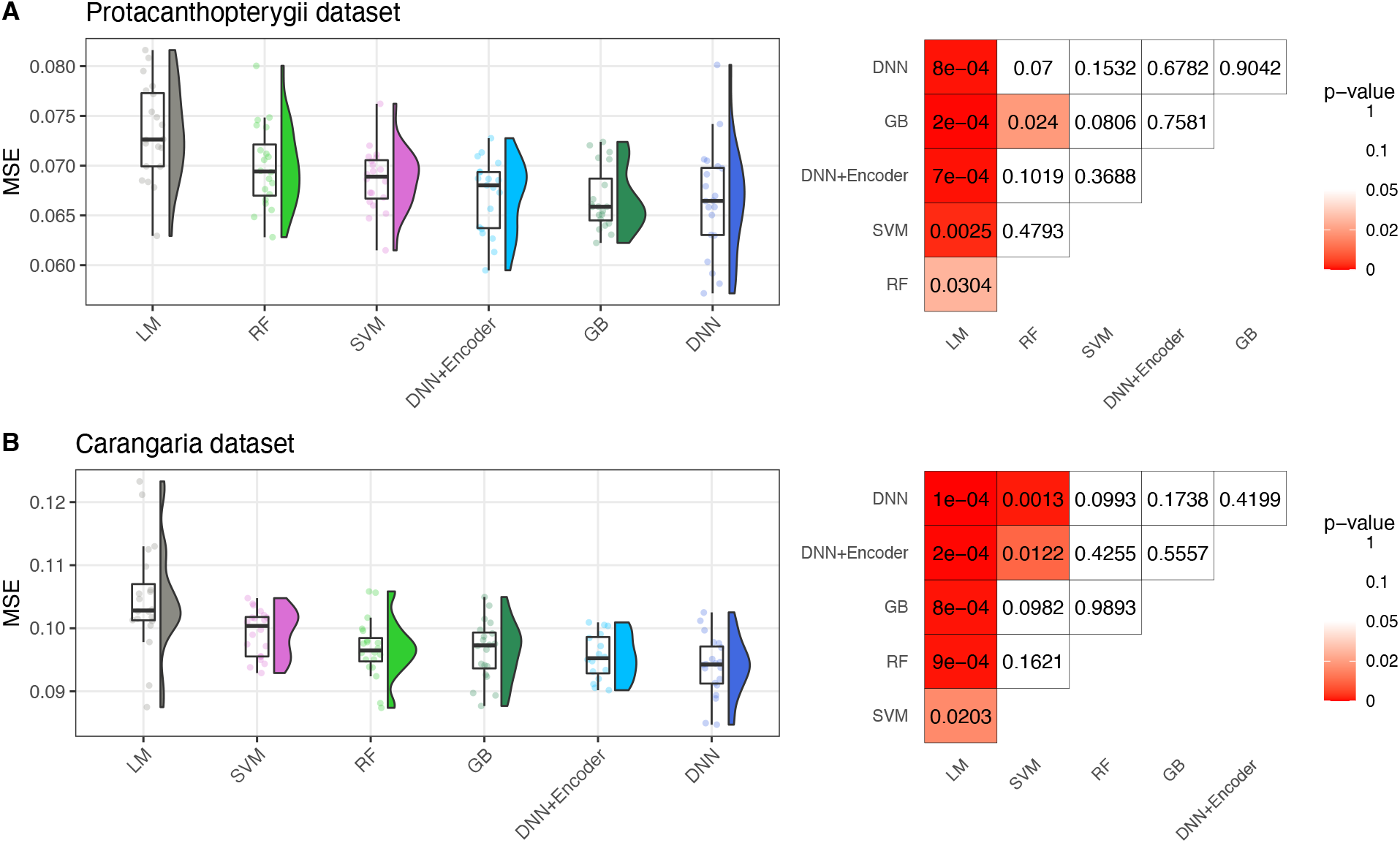
Testing MSE distribution from 20 rounds of CV for each model, and pairwise comparison of model testing MSE based on Wilcox Rank Sum Test. Panels **A)** and **B)** show the distribution of MSE and p-values for the Protacantopterygii and Carangaria datasets, respectively. Box plots are arranged in descending order of the average MSE for each model. LM: Linear Model; SVM: Support Vector Machine; RF: Random Forest; GB: Gradient Boosting; DNN: Deep Neural Network; and DNN + Encoder: Deep Neural Network with Encoder layers.

Tested features (see table S4) displayed different weights in each dataset when the DNN model predicts AU p-values. When we ordered the tested features as a function of the average increase of MSE from the DNN model + 40 rounds of CV (feature importance analysis, see Materials and Methods), the ordering of features from top to bottom was differed across datasets (Fig. 5). For instance, while the feature importance profile of Protacanthopterygii identifies the variance of gap percentage per sequence at codon position 3 (“gap_var_pos3”) as the top feature, this same feature ranked 32^nd^ in the feature importance profile of Carangaria. Similarly, the top feature from the feature importance profile of Carangaria dataset was the variance of GC-content per sequence at codon position 3 (“gc_mean_pos3”), but this feature ranked 7^th^ in the feature importance profile of Protacanthopterygii. The top features for both datasets were sequence-based features. In contrast to the Carangaria dataset, the Protacanthopterygii dataset did not have a clearly leading feature; the fold difference between “gap_var_pos3” and its second top feature “LB_std” (standard deviation of the long branch score per sequence as described by Struck 2014) was 1.0009. In contrast, for the Carangaria dataset the fold difference between “gc_mean_pos3” and its second top feature “inter_len_mean” (mean of internal branch lengths in the tree) was 2.8132.

**Figure 5.**
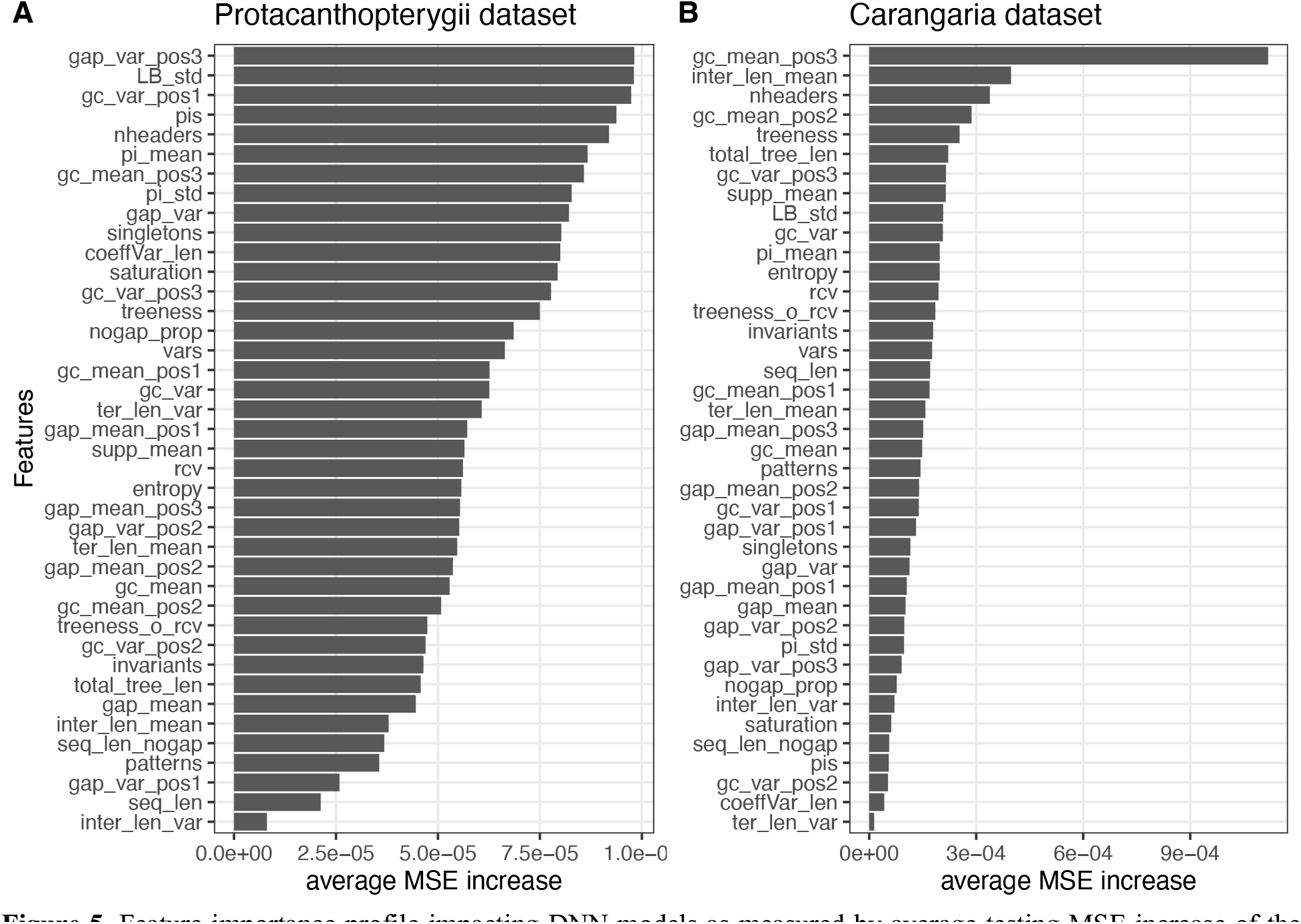
Feature importance profile impacting DNN models as measured by average testing MSE increase of the model. **A)** Protacanthopterygii dataset; **B)** Carangaria dataset. Bars are positioned in descending order. The top two features for the Protacantopterygii dataset were the variance of gap percentage per sequence at codon position 3 (“gap_var_pos3”) and the standard deviation of the long branch score per sequence (“LB_std”, as described by Struck 2014). The top two features for the Carangaria dataset were the mean of GC content per sequence at codon position 3 (“gc_mean_pos3”) and the mean of internal branch lengths in the tree (“inter_len_mean”). The remaining features have a relatively lower impact on the DNN model (as long as they are away from the top position on the y-axis). Table S4 shows both complete names and descriptions of tested features.

## 4. Discussion

### 4.1 Phylogenomic analyses

We generated a new dataset for Protacanthopterygii and utilized an existing dataset for Carangaria to estimate phylogenetic trees through concatenation-based maximum likelihood (ML) and multispecies coalescent (MSC) methods. As a result, two major hypotheses emerged for each dataset. In this section, we will focus the discussion on phylogenetic relationships within Protacanthopterygii. For a detailed discussion on the phylogeny of Carangaria and the monophyly of flatfishes, please refer to Duarte-Ribeiro et al. (2023, under review).

The phylogeny of Protacanthopterygii, considered as the sister group of Neoteleostei and the early diverging Euteleostei (Greenwood 1966), has been the subject of multiple hypotheses based on morphology and molecular data, including the present study (Fig. 3). While there is consensus among previous studies regarding two distinct and consistent supra-ordinal clades (Esociformes + Salmoniformes and Stomiiatiformes + Osmeriformes) (e.g., see Li et al., 2010; Near et al. 2012; Campbell et al., 2013; Betancur-R. et al., 2017), disagreements persist in other aspects. Specifically, authors have not reached a consensus on the composition and monophyly of Protacanthopterygii (Campbell et al., 2013; Miya and Nishida, 2015; Mirande et al., 2017; Betancur-R. et al., 2017). For instance, the most recent edition of “Fishes of the World” (Nelson et al., 2016) limits Protacanthopterygii to Salmoniformes and Esociformes. However, other molecular-based phylogenetic classifications (Betancur-R. et al., 2013, 2017) include Argentiniformes, Galaxiiformes, Salmoniformes, Esociformes, Osmeriformes, and Stomiatiformes (Stomiati) within Protacanthopterygii, with Osmeriformes and Stomiatiformes being the sister taxon to Neoteleostei. Similarly, our study also fails to resolve Protacanthopterygii (*sensu* Greenwood 1966) as a monophyletic group and sister clade of Neoteleostei. Given the lack of consensus among studies regarding the limits and monophyly of Protacanthopterygii as a sister clade of Neoteleostei and considering that the pre-cladistic establishment of Protacanthopterygii predates the acceptance of current phylogenetic methodology and interpretation, it can be argued that the delimitation of Protacanthopterygii *sensu* Greenwood 1966 as a natural group is primarily be due to historical reasons. Rather than insisting on identification of taxa that belong to this group, it may be more productive to understand the implications of the branching pattern among these taxa along the backbone of the fish Tree of Life as depicted in Figure 3a (see below).

### 4.2 Machine Learning and Feature Importance

By conducting Feature Importance analysis (see Material and Method section), we successfully identified a set of factors that contribute to uncertainty within each dataset. It is worth mentioning that these factors are unique to their respective datasets. Our analyses employ AU p-values, which have been widely recognized as a valuable tool for assessing the data signal among a range of tree options (Arcila et al. 2017). It takes into account the distribution of site-log likelihoods through a hypothesis-testing framework (Efron, Halloran, and Holmes 1996; Shimodaira 2002) and can be utilized as a label in machine learning models. If the AU p-value is significantly low, it provides strong evidence in favor of the alternative hypothesis (i.e., either H1 or H2 topology), based on the bootstrap distribution derived repeatedly from the log-likelihood data. This is particularly beneficial when working with binary classification models. However, the task of selecting between hypotheses using the AU p-value and its corresponding threshold can be challenging since certain trees may receive equal support from the available data (Simion et al. 2020). In our study, we encountered such cases, which prompted us to employ a multioutput regression model to predict both p-values as labels instead of utilizing a single p-value in a classification mode. This approach is more suitable in situations where datasets provide evidence for both hypotheses rather than favoring one hypothesis over the other.

The poor performance of the linear regression model compared to all non-linear models, with no noticeable difference observed among the non-linear models (Fig. 4), indicates that the feature space is highly non-linear. This is a characteristic that is increasingly being recognized as common in phylogenetic datasets (Billera, Holmes and Vogtmann 2001; Zou et al., 2020; Jiang et al. 2022). In contrast, models that incorporate non-linear interactions of factors provide an alternative approach to previous studies, which focused on identifying factors underlying phylogenetic uncertainty without assessing their non-linear interactions (Shen et al. 2020, Vakan et al. 2021, Duchêne et al. 2021). Convex algorithms such as SVM, which assume that local optima are also global optima, performed better than the linear model in both datasets, and they do not require a large amount of data. However, non-convex algorithms such as decision tree-based methods (e.g., RF, GB) and neural network-based methods (e.g., DNN, DNN+Encoder) utilize more instances to thoroughly search the feature space, as they can handle multiple local minima (Hastie et al., 2009; LeCun et al. 2015). Since our datasets are of medium size, with around 1000 instances each, these non-convex algorithms have the advantage of exploring the feature space more comprehensively. With the increasing availability of chromosome-level genomes (Simakov et al., 2022; Schultz et al., 2023), it is expected that DNNs will capture even more intricate non-linear relationships in whole-genome phylogenomics.

The DNN model that achieved the lowest testing MSE used various regularization techniques, including early stopping and dropout, which enhanced its generalizability as a non-linear model. Early stopping halts the training process when the MSE of the testing set starts increasing, while dropout randomly deactivates neurons during training, enabling the network to learn from a more extensive set of neurons after many iterations (Srivastava et al. 2014). We obtained the optimal dropout rate for each dataset through hyperparameter tuning. Additionally, we discovered that each dataset had a unique optimal dropout rate as well as other hyperparameters (see *Supplementary Data*). Since previous studies have shown significant accuracy improvements by including a hyperparameter tuning step (Zou and Hastie 2005, Krizhevsky et al. 2017), we recommend including hyperparameter tuning as an important component of the machine learning model training process in phylogenetic-based studies. Although the DNN is a generalizable model, the explicit interaction of features remains unknown. Therefore, we conducted feature importance analyses.

The significance of the features obtained is dataset-specific; hence, the signal varies among datasets. For example, in the Protacanthopterygii dataset, the variance of gap percentage per sequence at codon position 3 (represented as “gap_var_pos3” in Fig. 5A) emerged as the top feature. This observation can be attributed to the fact that the considered lineages in this dataset diverged at least 100 million years ago (Betancur-R. et al., 2017; Hughes et al. 2018), and a high incidence of indels is expected, particularly across different taxonomic scales, an issue that is further compounded by the diversified sampling strategy applied for building the Protacanthopterygii dataset. Notably, nodal support for order-level clades was strong for both hypotheses (Fig. 3). However, the indel variance within individual gene alignments can introduce errors in gene tree estimation, potentially leading to statistical inconsistencies in tree inference using ASTRAL (H2 of the Protacanthopterygii dataset) (Molloy and Warnow 2018). Furthermore, variation in root-to-tip distances, as represented by “LB_std” (Fig. 5A), has also been associated with statistical inconsistency in multispecies coalescent tree inference (Vankan, Ho and Duchêne 2022). To mitigate potential biases introduced by these factors, two possible solutions can be considered. First, increasing the number of species would lead to more accurate gene trees through a denser alignment (Xi et al. 2016). Alternatively, increasing the number of sites, for example by sampling sliding windows from whole genome alignments, can provide the opportunity to examine genomic regions directly affecting the branches of interest (e.g., the rogue placement of Argentiniformes). It has been observed that using this approach in large-scale datasets can improve the signal-to-noise ratio (Zhang et al. 2021; Di Franco et al. 2022), assuming there is no DNA sequence evolution model misspecification (Kumar et al. 2012) or introgression (Solís-Lemus et al. 2016). For the time being, our study provides justification to support H1, the topology based on concatenation-based ML (Fig 3).

Analyses of the Carangaria dataset revealed that the GC content at third codon positions (‘gc_mean_pos3’) was the most significant feature, consistent with previous findings identifying GC content as a biasing factor in this group of species (Betancur-R. et al., 2013). In cases where groups undergo rapid radiations early in their history, it becomes necessary to incorporate a heterogeneous substitution process that accounts for shifting rates within sites over time (Tuffley and Steel 1998; Hulsenbeck 2002; Jermiin et al., 2017, 2020). This is due to the varying probabilities of state changes within the rapidly radiating group compared to the rest of the tree, a phenomenon known as heterotachy (Lopez et al., 2002; Kolaczkowski and Thornton 2004; Kapli et al., 2020). While the majority of phylogenetic studies assume homogeneity of substitution rates across the entire tree to reduce computational burden (Pupko and Mayrose 2020), this assumption may lead to inaccurate phylogenetic reconstruction when the dataset exhibits bias towards certain states, such as GC content, within a specific subset of species (Hulsenbeck 2002; Kolaczkowski and Thornton 2004; Romiguier and Roux 2017; Baele et al. 2021). The monophyletic status of flatfishes has been supported by analyses with a small set of exon markers prone to GC biases (Fig. S1, H1 in the Flatfishes dataset; Betancur-R. et al., 2013), but other studies (Campbell et al. 2013; Lü et al. 2021) suggest flatfishes are polyphyletic (Fig S1, H2 in the Flatfishes dataset), implying dual evolutionary origins of their asymmetric body plan. However, when GC biases are explicitly modeled, using for example non-homogeneous GHOST model (Crotty et al., 2020), then analyses of exon markers largely support the hypothesis of flatfish monophyly (Duarte-Ribeiro et al., under review), albeit with a reduced taxonomic dataset due to computational intractability of the GHOST model. This result is also consistent with other datasets using UCE markers (which are less affected by GC biases; Jarvis et al. 2014) analyzed using both homogenous (Harrington et al. 2016) and non-homogeneous (Duarte-Ribeiro et al., under review) models.

By gaining an understanding of the underlying factors that influence a dataset, systematists can make more informed decisions about the appropriate type of phylogenetic analysis or data pre-processing methods to employ before inputting the data into existing software for phylogenetic inference. However, if we let the machine learning models such as Convolutional Neural Networks (Fonseca et al. 2021; Yang et al., 2022) or Graph Neural Networks (Voznica et al., 2022; Lajaaiti et al., 2023) learn optimized features directly from the alignment or tree during the training process, we may sacrifice model explainability, which is a current challenge in applying machine learning models to biological datasets (Sapoval et al., 2022). This is because the features learned from the model may not correlate with existing literature that addresses factors influencing phylogenetic uncertainty. In contrast, our study’s Feature Importance analyses demonstrate clear connections with previous studies, reinforcing the notion that well-defined features can inform decision-making in heuristic search moves within the tree space (Azouri et al., 2021, 2023) and predict the difficulty of a dataset for phylogenetic reconstruction (Haag et al., 2022). However, our study also suggests, particularly after analyzing the Carangaria dataset, that current machine learning models in phylogenetics that use empirical datasets should be trained with data that is agnostic to phylogenetic model assumptions such as homogeneity, stationarity, and reversibility (Pupko and Mayrose, 2020).

It has been observed that non-parametric bootstrap p-values, which serve as the statistical basis for labeling our machine learning models, do not converge to a single point value as the number of sites approaches infinity (Susko, 2009; Huang et al., 2020). Additionally, depending on the curvature of the tree space, selection bias can arise in the calculations of p-values (Efron, 1996; Susko, 2009). While previous studies have addressed this problem (Shimodaira 2002; Anisimova and Gascuel 2006), including the avoidance of p-values altogether (Minh et al., 2020), and our sequences are relatively short, with an average length of approximately 400 bp, we recognize the importance of further studies to address this issue in the context of machine learning labels. For instance, exploring the use of more conservative hypothesis-testing tools such as the KH test (Kishino and Hasegawa, 1989) to label the machine learning model could be considered when dealing with longer sequences. While the random permutation approach was effective in assessing factor interactions, more theoretically sounded methods like SHAP values (Lundberg et al., 2020) can provide comprehensive marginal contributions of feature importance. Although Dropout demonstrated effectiveness in our study, we acknowledge that incorporating additional regularization techniques, such as L2 norm in combination with Dropout, could further improve model generalization (Srivastava et al. 2014). Lastly, to enhance training efficiency for large datasets, transferring base layers that learned low-level features from pre-trained models into a new model can be beneficial, as the performance difference between DNN and DNN+Encoder was not significant (Fig. 4).

### 4.3 Conclusions

This study introduces a novel approach that explores factors that can explain differences among competing phylogenetic hypotheses by utilizing machine learning models to capture non-linear interactions between such factors. The obtained insights can greatly enhance decision-making in phylogenetic analyses, aiding in the selection of appropriate DNA sequence models and data transformation methods. While machine learning has demonstrated its utility in various aspects of phylogenetic analysis, such as biogeography models (Smith et al., 2017; Fonseca et al., 2021), phylodynamic parameter estimation (Voznica et al., 2022), ancestral state reconstruction (Theobald et al., 2022), and improving heuristics for phylogenetic inference (Azouri et al., 2021, 2023), the quality of these models ultimately depends on the quality of the training data, which can inherently carry its own uncertainty, as we showed in this study. Therefore, the pursuit of identifying factors that influence the uncertainty in phylogenomic datasets and developing models that effectively capture non-linear relationships among features in phylogenetic inference is essential for advancing our understanding of the tree of life.

Our analysis of the two phylogenomic datasets reveals that distinct dataset-specific factors contribute unequally to phylogenetic uncertainty. Considering the significance of the top features in the Protacanthopterygii dataset, which are associated with errors in gene tree estimations, we suggest increasing the number of species and sites in this dataset. Nevertheless, this result contributes to the ongoing resolution of the debated early evolutionary history of teleost fishes (Betancur et al., 2017; Hughes et al., 2018). While there is phylogenetic evidence supporting flatfish monophyly based on molecular data (i.e., the H1 hypothesis in the Carangaria dataset, Fig. S1), additional lines of evidence, such as morphology and developmental pathways (Duarte-Ribeiro et al., under review), can confirm this trend and provide further exploration of this challenging group undergoing rapid radiation.

## Acknowledgements

This project was funded by the National Science Foundation (NSF) [DEB-1932759 and DEB-2225130 to R.B., DEB-2015404 and DEB-2144325 to D.A., and DEB-1541554 to G.O.]. Computationally demanding extraction of feature information from thousands of gene trees and DNA alignments was done using resources provided by the OSG Consortium (OSG 2006; Pordes et al. 2007; Sfiligoi et al. 2009), which is supported by the NSF awards #2030508 and #1836650.

## Data availability

Read sequence information was deposited into NCBI under the following Bioproject Accession ID: PRJNA1013549. The optimal hyperparameters for each model, as well as the phylogenetic trees and alignments used for the present study, can be found here: https://github.com/Ulises-Rosas/dissectingFactors.

## Code availability

The code for hyperparameter tuning, model fitting, cross-validations, and feature importance analyses (specifically for Deep Neural Networks) for all the supervised machine learning models tested in this study can be found at the following repository: https://github.com/Ulises-Rosas/dissectingFactors. This repository also contains the datasets that were tested. Additionally, the Python-based command-line interface for quality control is accessible at https://github.com/Ulises-Rosas/fishlifeqc, and the Python-based command-line for GGI analyses and dataset features extraction can be found at https://github.com/Ulises-Rosas/GGpy.

