## Supplementary Material for "Dissecting Factors Underlying Phylogenetic Uncertainty Using Machine Learning Models"

### S.1 Quality Control

We used a quality control (QC) pipeline to address potential sources of contamination, mislabeling, sequencing errors, or other confounding features of the data. All steps involved in this pipeline were integrated into a Python-based command-line interface called `"fishlifeqc"` (available at: github.com/ulises-rosas/fishlifeqc). We based this pipeline on the conceptual framework proposed by Simion et al. (2017); Arcila et al. (2021) and it currently includes three main functions:

i Missing data: Missing data has well known impact on the phylogeny estimation (Xi et al., 2016; Smirnov and Warnow, 2021; Portik and Wiens, 2021), and we included a step for controlling the amount of missing data in this pipeline. Since adoption of a universal threshold for eliminating poorly aligned regions can be problematic (Molloy and Warnow, 2018), our implementation eliminates complete exon alignments with less than 75% taxa. After this step, we excluded 79 and 114 exon alignments for the Protacanthopterygii and Carangaria datasets, respectively. A complete list of options for this step can be obtained with the command "`fishlifeqc mdata -h`".

ii Mislabeling: Errors involving mislabeling of samples (sequences) can induce phylogenetic errors or misleading conclusions on, for instance, the timing of coalescent events (Song et al., 2012; Springer and Gatesy, 2016). This step is designed to verify correct species identification in a dataset by using mitochondrial COI barcode data by taking leveraging the BOLD ID Engine webservice. After this step, we update 2 and 15 species names for the Protacanthopterygii and Carangaria datasets, respectively. In the case of the Carangaria dataset, we eliminated 20 samples as they belong to non-Carangarian species. A complete list of options for this step can be obtained with the command "`fishlifeqc bold -h`".

iii Taxon misplacement due to orthology error or contamination: This step compares the terminal branch lengths between a reference tree and the gene tree for each exon alignment, and it was first proposed by Simion et al. (2017, 2020) as their "branch length correlation" or BLC test. We removed sequences for which the branch-length ratio $> 5$ and we used the Pearson correlation coefficient ($R^2$) between branch lengths in the same two trees to identify (and eliminate) gene alignments that are outliers. After this step, we excluded 30 exon alignments for the Protacanthopterygii dataset. A complete list of options for this step can be obtained with the command "`fishlifeqc bl -h`".

Table S1: List of species examined of the Carangaria dataset

| **Family** | **Genera** | **Species** | **Accession** | **Data type** |
|---|---|---|---|---|
| Acanthuridae | | | | |
| | Acanthurus | | | |
| | | *Acanthurus mata* | Duarte-Ribeiro et al. (USNMAG9RD41) | Genome |
| Achiridae | | | | |
| | Achirus | | | |
| | | *Achirus lineatus* | Duarte-Ribeiro et al. (KU5115) | Genome |
| | Catathyridium | | | |
| | | *Catathyridium jenynsii* | Duarte-Ribeiro et al. (UNMDP539) | Genome |
| | Gymnachirus | | | |
| | | *Gymnachirus melas* | Duarte-Ribeiro et al. (KU3932) | Genome |
| | Trinectes | | | |
| | | *Trinectes fluviatilis* | Duarte-Ribeiro et al. (STRIBFT04923) | Genome |
| | | *Trinectes fluviatilis* | Duarte-Ribeiro et al. (UMSNH12340) | Genome |
| | | *Trinectes fonsecensis* | Duarte-Ribeiro et al. (UMSNH8485) | Genome |
| | | *Trinectes inscriptus* | Duarte-Ribeiro et al. (USNMAC8CC29) | Genome |
| | | *Trinectes maculatus* | Duarte-Ribeiro et al. (KU1501) | Genome |
| | | *Trinectes sp* | Duarte-Ribeiro et al. (UMSNH16683) | Genome |
| Ambassidae | | | | |
| | Ambassis | | | |
| | | *Ambassis nalua* | Duarte-Ribeiro et al. (EPLATE_42_G04) | Genome |
| Apogonidae | | | | |
| | Apogon | | | |
| | | *Apogon robinsi* | Duarte-Ribeiro et al. (EPLATE_06_F08) | Genome |
| Batrachoididae | | | | |
| | Opsanus | | | |
| | | *Opsanus tau* | Duarte-Ribeiro et al. (EPLATE_23_F09) | Genome |
| Blenniidae | | | | |
| | Exallias | | | |
| | | *Exallias brevis* | Duarte-Ribeiro et al. (EPLATE_42_A11) | Genome |

*Continued on next page*

Table S1 – *Continued from previous page*

| **Family** | **Genera** | **Species** | **Accession** | **Data type** |
|---|---|---|---|---|
| Bothidae | | | | |
| | Arnoglossus | | | |
| | | *Arnoglossus aspilos* | Duarte-Ribeiro et al. (CSIROGT668) | Genome |
| | | *Arnoglossus macrolophus* | Duarte-Ribeiro et al. (CSIROGT671) | Genome |
| | | *Arnoglossus waitei* | Duarte-Ribeiro et al. (CSIROGT7205) | Genome |
| | Asterorhombus | | | |
| | | *Asterorhombus cocosensis* | Duarte-Ribeiro et al. (KU7102) | Genome |
| | | *Asterorhombus filifer* | Duarte-Ribeiro et al. (USNMAG9RT70) | Genome |
| | Bothus | | | |
| | | *Bothus lunatus* | Duarte-Ribeiro et al. (USNMT154) | Genome |
| | | *Bothus maculiferus* | Duarte-Ribeiro et al. (USNMAC4YY74) | Genome |
| | | *Bothus mancus* | Duarte-Ribeiro et al. (UMSNH34382) | Genome |
| | | *Bothus ocellatus* | Duarte-Ribeiro et al. (UMSNH29267) | Genome |
| | | *Bothus pantherinus* | Duarte-Ribeiro et al. (KU5642) | Genome |
| | | *Bothus podas* | Duarte-Ribeiro et al. (USNMAD9NE07) | Genome |
| | | *Bothus robinsi* | Duarte-Ribeiro et al. (KU1169) | Genome |
| | | Bothus sp1 | Duarte-Ribeiro et al. (CSIROGT7201) | Genome |
| | Crossorhombus | | | |
| | | *Crossorhombus kobensis* | Duarte-Ribeiro et al. (KU2489) | Genome |
| | Engyophrys | | | |
| | | *Engyophrys sanctilaurentii* | Duarte-Ribeiro et al. (USNMAG7PC76) | Genome |
| | | *Engyophrys senta* | Duarte-Ribeiro et al. (KU5128) | Genome |
| | Engyprosopon | | | |
| | | *Engyprosopon bleekeri* | Duarte-Ribeiro et al. (CSIROGT7231) | Genome |
| | | *Engyprosopon maldivense* | Duarte-Ribeiro et al. (CSIROGT7237) | Genome |
| | | *Engyprosopon multisquama* | Duarte-Ribeiro et al. (KU2503) | Genome |
| | Grammatobothus | | | |
| | | *Grammatobothus pennatus* | Duarte-Ribeiro et al. (CSIROGT7186) | Genome |
| | Japonolaeops | | | |
| | | *Japonolaeops dentatus* | Duarte-Ribeiro et al. (CSIROGT659) | Genome |
| | Laeops | | | |
| | | *Laeops kitaharae* | Duarte-Ribeiro et al. (KU2506) | Genome |
| | Lophonectes | | | |
| | | *Lophonectes gallus* | Duarte-Ribeiro et al. (CSIROGT447) | Genome |
| | Monolene | | | |
| | | *Monolene asaedae* | Duarte-Ribeiro et al. (USNMAG7PH93) | Genome |
| | | *Monolene maculipinna* | Duarte-Ribeiro et al. (USNMAG7PD52) | Genome |
| | Psettina | | | |
| | | *Psettina iijimae* | Duarte-Ribeiro et al. (KU2511) | Genome |
| | | *Psettina senta* | Duarte-Ribeiro et al. (CSIROGT674) | Genome |
| | | Psettina sp1 | Duarte-Ribeiro et al. (CSIROGT7164) | Genome |
| | Trichopsetta | | | |
| | | *Trichopsetta ventralis* | Duarte-Ribeiro et al. (KU5085) | Genome |
| Carangidae | | | | |
| | Alectis | | | |
| | | *Alectis alexandrina* | Duarte-Ribeiro et al. (ODU1437) | Genome |
| | | *Alectis ciliaris* | Duarte-Ribeiro et al. (UMSNH16987) | Genome |
| | | *Alectis indica* | Duarte-Ribeiro et al. (USNMT8966) | Genome |
| | | *Alectis indica* | Duarte-Ribeiro et al. (USNMT8967) | Genome |
| | Alepes | | | |

*Continued on next page*

Table S1 – *Continued from previous page*

| Family | Genera | Species | Accession | Data type |
|---|---|---|---|---|
| | | *Alepes djedaba* | Duarte-Ribeiro et al. (USNMT8985) | Genome |
| | | *Alepes kleinii* | Duarte-Ribeiro et al. (USNMT8986) | Genome |
| | | *Alepes vari* | Duarte-Ribeiro et al. (USNMAC1VJ47) | Genome |
| | Atule | | | |
| | | *Atule mate* | Duarte-Ribeiro et al. (KU7225) | Genome |
| | Campogramma | | | |
| | | *Campogramma glaycos* | Duarte-Ribeiro et al. (ODU1311) | Genome |
| | Carangoides | | | |
| | | *Carangoides armatus* | Duarte-Ribeiro et al. (USNMAB6QR16) | Genome |
| | | *Carangoides bajad* | Duarte-Ribeiro et al. (USNMAG9RE46) | Genome |
| | | *Carangoides bartholomaei* | Duarte-Ribeiro et al. (FL1649) | Genome |
| | | *Carangoides dinema* | Duarte-Ribeiro et al. (USNMAG9RE59) | Genome |
| | | *Carangoides equula* | Duarte-Ribeiro et al. (ODU1315) | Genome |
| | | *Carangoides ferdau* | Duarte-Ribeiro et al. (KU6972) | Genome |
| | | *Carangoides fulvoguttatus* | Duarte-Ribeiro et al. (USNMT6973) | Genome |
| | | *Carangoides gymnostethus* | Duarte-Ribeiro et al. (KU6775) | Genome |
| | | *Carangoides hedlandensis* | Duarte-Ribeiro et al. (ODU4223) | Genome |
| | | *Carangoides hedlandensis* | Duarte-Ribeiro et al. (USNMAG9RD31) | Genome |
| | | *Carangoides humerosus* | Duarte-Ribeiro et al. (CSIROGT4745) | Genome |
| | | *Carangoides malabaricus* | Duarte-Ribeiro et al. (ODU1320) | Genome |
| | | *Carangoides oblongus* | Duarte-Ribeiro et al. (FMNH126718) | Genome |
| | | *Carangoides orthogrammus* | Duarte-Ribeiro et al. (UMSNH35087) | Genome |
| | | *Carangoides otrynter* | Duarte-Ribeiro et al. (UMSNH11087) | Genome |
| | | *Carangoides plagiotaenia* | Duarte-Ribeiro et al. (KU7154) | Genome |
| | | *Carangoides praeustus* | Duarte-Ribeiro et al. (USNMAG9RF82) | Genome |
| | | *Carangoides talamparoides* | Duarte-Ribeiro et al. (USNMAG9RF18) | Genome |
| | | *Carangoides vinctus* | Duarte-Ribeiro et al. (UMSNH20576) | Genome |
| | Caranx | | | |
| | | *Caranx bucculentus* | Duarte-Ribeiro et al. (CSIRO2Cbuc) | Genome |
| | | *Caranx caballus* | Duarte-Ribeiro et al. (FL1226) | Genome |
| | | *Caranx caninus* | Duarte-Ribeiro et al. (UMSNH24963) | Genome |
| | | *Caranx hippos* | Duarte-Ribeiro et al. (UMSNH16902) | Genome |
| | | *Caranx hippos* | Duarte-Ribeiro et al. (UPRFL1629) | Genome |
| | | *Caranx ignobilis* | Duarte-Ribeiro et al. (KU4947) | Genome |
| | | *Caranx latus* | Duarte-Ribeiro et al. (UMSNH16645) | Genome |
| | | *Caranx lugubris* | Duarte-Ribeiro et al. (UMSNH34515) | Genome |
| | | *Caranx papuensis* | Duarte-Ribeiro et al. (USNMAG9RT90) | Genome |
| | | *Caranx rhonchus* | Duarte-Ribeiro et al. (ODU0849) | Genome |
| | | *Caranx ruber* | Duarte-Ribeiro et al. (UMSNH14130) | Genome |
| | | *Caranx ruber* | Duarte-Ribeiro et al. (UPRFL0243) | Genome |
| | | *Caranx senegallus* | Duarte-Ribeiro et al. (ODU2506) | Genome |
| | | *Caranx sexfasciatus* | Duarte-Ribeiro et al. (KU6826) | Genome |
| | | *Caranx sexfasciatus* | Duarte-Ribeiro et al. (UMSNH24909) | Genome |
| | | *Caranx sexfasciatus* | Duarte-Ribeiro et al. (UMSNH24930) | Genome |
| | | *Caranx tille* | Duarte-Ribeiro et al. (ODU1330) | Genome |
| | Chloroscombrus | | | |
| | | *Chloroscombrus chrysurus* | Duarte-Ribeiro et al. (UMSNH29728) | Genome |
| | | *Chloroscombrus orqueta* | Duarte-Ribeiro et al. (UPRFL1337) | Genome |
| | Decapterus | | | |

*Continued on next page*

Table S1 – *Continued from previous page*

| Family | Genera | Species | Accession | Data type |
|---|---|---|---|---|
| | | *Decapterus Selar* | Duarte-Ribeiro et al. (crumenophthalmus_FL1643) | Genome |
| | | *Decapterus cf* | Duarte-Ribeiro et al. (macrosoma_FL1335) | Genome |
| | | *Decapterus kurroides* | Duarte-Ribeiro et al. (USNMAG9RF06) | Genome |
| | | *Decapterus macarellus* | Duarte-Ribeiro et al. (KU191) | Genome |
| | | *Decapterus muroadsi* | Duarte-Ribeiro et al. (USNMT8984) | Genome |
| | | *Decapterus punctatus* | Duarte-Ribeiro et al. (UMSNH33701) | Genome |
| | Elagatis | | | |
| | | *Elagatis bipinnulata* | Duarte-Ribeiro et al. (KU6820) | Genome |
| | | *Elagatis bipinnulata* | Duarte-Ribeiro et al. (UMSNH11021) | Genome |
| | Hemicaranx | | | |
| | | *Hemicaranx amblyrhynchus* | Duarte-Ribeiro et al. (KU5142) | Genome |
| | | *Hemicaranx leucurus* | Duarte-Ribeiro et al. (UMSNH22493) | Genome |
| | | *Hemicaranx leucurus* | Duarte-Ribeiro et al. (UPRFL1510) | Genome |
| | Lichia | | | |
| | | *Lichia amia* | Duarte-Ribeiro et al. (ODU1342) | Genome |
| | Megalaspis | | | |
| | | *Megalaspis cordyla* | Duarte-Ribeiro et al. (KU4705) | Genome |
| | Naucrates | | | |
| | | *Naucrates ductor* | Duarte-Ribeiro et al. (USNMAB6QQ88) | Genome |
| | Oligoplites | | | |
| | | *Oligoplites altus* | Duarte-Ribeiro et al. (UMSNH10116) | Genome |
| | | *Oligoplites altus* | Duarte-Ribeiro et al. (UMSNH12517) | Genome |
| | | *Oligoplites saliens* | Duarte-Ribeiro et al. (UNMDP1504) | Genome |
| | | *Oligoplites saurus* | Duarte-Ribeiro et al. (UPRFL0970) | Genome |
| | Parastromateus | | | |
| | | *Parastromateus niger* | Duarte-Ribeiro et al. (USNMAG9RD57) | Genome |
| | Parona | | | |
| | | *Parona signata* | Duarte-Ribeiro et al. (UNMDPT0431) | Genome |
| | Pseudocaranx | | | |
| | | *Pseudocaranx chilensis* | Duarte-Ribeiro et al. (ODU1355) | Genome |
| | | *Pseudocaranx dentex* | Duarte-Ribeiro et al. (USNMAD9NF93) | Genome |
| | | *Pseudocaranx georgianus* | Duarte-Ribeiro et al. (ODU2288) | Genome |
| | | *Pseudocaranx wrighti* | Duarte-Ribeiro et al. (CSIRO1Pwri) | Genome |
| | Scomberoides | | | |
| | | *Scomberoides lysan* | Duarte-Ribeiro et al. (USNMT5653) | Genome |
| | | *Scomberoides tala* | Duarte-Ribeiro et al. (CSIROIN01832) | Genome |
| | | *Scomberoides tol* | Duarte-Ribeiro et al. (USNMAG9RF16) | Genome |
| | Selar | | | |
| | | *Selar crumenophthalmus* | Duarte-Ribeiro et al. (FL1338) | Genome |
| | | *Selar crumenophthalmus* | Duarte-Ribeiro et al. (UMSNH12296) | Genome |
| | Selene | | | |
| | | *Selene brevoortii* | Duarte-Ribeiro et al. (KU8505) | Genome |
| | | *Selene brevoortii* | Duarte-Ribeiro et al. (UMSNH11513) | Genome |
| | | *Selene dorsalis* | Duarte-Ribeiro et al. (ODU1369) | Genome |
| | | *Selene orstedii* | Duarte-Ribeiro et al. (STRIBFT00327) | Genome |
| | | *Selene peruviana* | Duarte-Ribeiro et al. (STRIBFT10081) | Genome |
| | | *Selene setapinnis* | Duarte-Ribeiro et al. (UMSNH29808) | Genome |
| | | *Selene vomer* | Duarte-Ribeiro et al. (UMSNH29804) | Genome |

*Continued on next page*

Table S1 – *Continued from previous page*

| Family | Genera | Species | Accession | Data type |
|---|---|---|---|---|
| | Seriola | | | |
| | | *Seriola carpenteri* | Duarte-Ribeiro et al. (ODU1439) | Genome |
| | | *Seriola dumerili* | Duarte-Ribeiro et al. (KU5170) | Genome |
| | | *Seriola fasciata* | Duarte-Ribeiro et al. (KU5323) | Genome |
| | | *Seriola hippos* | Duarte-Ribeiro et al. (CSIRO2Ship) | Genome |
| | | *Seriola lalandi* | Duarte-Ribeiro et al. (UMSNH31308) | Genome |
| | | *Seriola peruana* | Duarte-Ribeiro et al. (UMSNH8407) | Genome |
| | | *Seriola rivoliana* | Duarte-Ribeiro et al. (UMSNH25769) | Genome |
| | | *Seriola zonata* | Duarte-Ribeiro et al. (KU3522) | Genome |
| | Trachinotus | | | |
| | | *Trachinotus africanus* | Duarte-Ribeiro et al. (CSIROIN00305) | Genome |
| | | *Trachinotus anak* | Duarte-Ribeiro et al. (CSIRO2Tana) | Genome |
| | | *Trachinotus baillonii* | Duarte-Ribeiro et al. (USNMT6824) | Genome |
| | | *Trachinotus blochii* | Duarte-Ribeiro et al. (USNMT6793) | Genome |
| | | *Trachinotus carolinus* | Duarte-Ribeiro et al. (UNMDPT2943) | Genome |
| | | *Trachinotus falcatus* | Duarte-Ribeiro et al. (UMSNH30046) | Genome |
| | | *Trachinotus goodei* | Duarte-Ribeiro et al. (UMSNH28915) | Genome |
| | | *Trachinotus goreensis* | Duarte-Ribeiro et al. (ODU1473) | Genome |
| | | *Trachinotus kennedyi* | Duarte-Ribeiro et al. (UMSNH22990) | Genome |
| | | *Trachinotus mookalee* | Duarte-Ribeiro et al. (ODU1379) | Genome |
| | | *Trachinotus ovatus* | Duarte-Ribeiro et al. (ODU2512) | Genome |
| | | *Trachinotus paitensis* | Duarte-Ribeiro et al. (UMSNH18640) | Genome |
| | | *Trachinotus rhodopus* | Duarte-Ribeiro et al. (UMSNH10673) | Genome |
| | | *Trachinotus stilbe* | Duarte-Ribeiro et al. (UMSNH17685) | Genome |
| | | *Trachinotus teraia* | Duarte-Ribeiro et al. (ODU1475) | Genome |
| | Trachurus | | | |
| | | *Trachurus declivis* | Duarte-Ribeiro et al. (CSIRO10Tdec) | Genome |
| | | *Trachurus lathami* | Duarte-Ribeiro et al. (KU5082) | Genome |
| | | *Trachurus murphyi* | Duarte-Ribeiro et al. (CSIRO1Tmur) | Genome |
| | | *Trachurus novaezelandiae* | Duarte-Ribeiro et al. (CSIRO1Tnov) | Genome |
| | | *Trachurus symmetricus* | Duarte-Ribeiro et al. (USNMT9419) | Genome |
| | Uraspis | | | |
| | | *Uraspis secunda* | Duarte-Ribeiro et al. (KU3988) | Genome |
| | | *Uraspis uraspis* | Duarte-Ribeiro et al. (ODU1262) | Genome |
| Centropomidae | | | | |
| | Centropomus | | | |
| | | *Centropomus ensiferus* | Duarte-Ribeiro et al. (UMSNH16600) | Genome |
| | | *Centropomus medius* | Duarte-Ribeiro et al. (KU8499) | Genome |
| | | *Centropomus nigrescens* | Duarte-Ribeiro et al. (FL1515) | Genome |
| | | *Centropomus pectinatus* | Duarte-Ribeiro et al. (STRIBFT00370) | Genome |
| | | *Centropomus robalito* | Duarte-Ribeiro et al. (UMSNH11363) | Genome |
| | | *Centropomus undecimalis* | Duarte-Ribeiro et al. (KU37) | Genome |
| | | *Centropomus unionensis* | Duarte-Ribeiro et al. (UMSNH22762) | Genome |
| | | *Centropomus viridis* | Duarte-Ribeiro et al. (KU8522) | Genome |
| | | *Centropomus viridis* | Duarte-Ribeiro et al. (UMSNH24594) | Genome |
| | Lates | | | |
| | | *Lates calcarifer* | Duarte-Ribeiro et al. (USNMAG9RD36) | Genome |
| | | *Lates japonicus* | Duarte-Ribeiro et al. (USNMT10313) | Genome |
| | | *Lates niloticus* | Duarte-Ribeiro et al. (CSIRO1Lnil) | Genome |
| | | *Lates uwisara* | Duarte-Ribeiro et al. (CSIROGT159) | Genome |

*Continued on next page*

Table S1 – *Continued from previous page*

| Family | Genera | Species | Accession | Data type |
|---|---|---|---|---|
| Chiasmodontidae | | | | |
| | Kali | | | |
| | | *Kali macrura* | Duarte-Ribeiro et al. (USNMT2229) | Genome |
| Citharidae | | | | |
| | Citharoides | | | |
| | | *Citharoides macrolepis* | Duarte-Ribeiro et al. (KU2468) | Genome |
| | Lepidoblepharon | | | |
| | | *Lepidoblepharon ophthalmolepis* | Duarte-Ribeiro et al. (CSIROGT5890) | Genome |
| Coryphaenidae | | | | |
| | Coryphaena | | | |
| | | *Coryphaena hippurus* | Duarte-Ribeiro et al. (UMSNH8410) | Genome |
| Cyclopsettidae | | | | |
| | Citharichthys | | | |
| | | *Citharichthys arctifrons* | Duarte-Ribeiro et al. (KU1090) | Genome |
| | | *Citharichthys cornutus* | Duarte-Ribeiro et al. (KU5196) | Genome |
| | | *Citharichthys gilberti* | Duarte-Ribeiro et al. (UMSNH12338) | Genome |
| | | *Citharichthys platophrys* | Duarte-Ribeiro et al. (USNMAG7PF78) | Genome |
| | | *Citharichthys sordidus* | Duarte-Ribeiro et al. (KU3255) | Genome |
| | | *Citharichthys xanthostigma* | Duarte-Ribeiro et al. (USNMT9170) | Genome |
| | Cyclopsetta | | | |
| | | *Cyclopsetta querna* | Duarte-Ribeiro et al. (UMSNH11512) | Genome |
| | Etropus | | | |
| | | *Etropus crossotus* | Duarte-Ribeiro et al. (UMSNH8395) | Genome |
| | | *Etropus crossotus* | Duarte-Ribeiro et al. (UMSNH8396) | Genome |
| | | *Etropus microstomus* | Duarte-Ribeiro et al. (KU1506) | Genome |
| | Syacium | | | |
| | | *Syacium longidorsale* | Duarte-Ribeiro et al. (UMSNH11532) | Genome |
| | | *Syacium maculiferum* | Duarte-Ribeiro et al. (UMSNH11529) | Genome |
| | | *Syacium micrurum* | Duarte-Ribeiro et al. (KU2) | Genome |
| | | *Syacium micrurum* | Duarte-Ribeiro et al. (KU5200) | Genome |
| | | *Syacium papillosum* | Duarte-Ribeiro et al. (KU1167) | Genome |
| Cynoglossidae | | | | |
| | Cynoglossus | | | |
| | | *Cynoglossus arel* | Duarte-Ribeiro et al. (CSIRO3R) | Genome |
| | | *Cynoglossus arelcf2* | Duarte-Ribeiro et al. (CSIROHK09) | Genome |
| | | *Cynoglossus cfarel* | Duarte-Ribeiro et al. (CSIROIN01920) | Genome |
| | | *Cynoglossus cfcynoglossus* | Duarte-Ribeiro et al. (CSIROIN01684) | Genome |
| | | *Cynoglossus interruptus* | Duarte-Ribeiro et al. (KU2478) | Genome |
| | | *Cynoglossus lida* | Duarte-Ribeiro et al. (CSIROIN02205) | Genome |
| | | *Cynoglossus maculipinnis* | Duarte-Ribeiro et al. (CSIROGT1274) | Genome |
| | | *Cynoglossus maculipinnis* | Duarte-Ribeiro et al. (USNMAC1VJ18) | Genome |
| | | *Cynoglossus ogilbyi* | Duarte-Ribeiro et al. (CSIROGT1270) | Genome |
| | | Cynoglossus sp3 | Duarte-Ribeiro et al. (CSIROGT1280) | Genome |
| | | Cynoglossus sp6 | Duarte-Ribeiro et al. (CSIROGT1258) | Genome |
| | | Cynoglossus sp7 | Duarte-Ribeiro et al. (CSIROGT1249) | Genome |
| | | Cynoglossus sp8 | Duarte-Ribeiro et al. (CSIROIN02204) | Genome |
| | Paraplagusia | | | |
| | | *Paraplagusia bilineata* | Duarte-Ribeiro et al. (CSIRO1vPbil) | Genome |
| | | *Paraplagusia bilineata* | Duarte-Ribeiro et al. (CSIROIN02407) | Genome |

*Continued on next page*

Table S1 – *Continued from previous page*

| **Family** | **Genera** | **Species** | **Accession** | **Data type** |
|---|---|---|---|---|
| | | *Paraplagusia bilineata* | Duarte-Ribeiro et al. (ODU5166) | Genome |
| | | *Paraplagusia blochii* | Duarte-Ribeiro et al. (CSIROIN02207) | Genome |
| | | *Paraplagusia cfguttata* | Duarte-Ribeiro et al. (CSIROHK18) | Genome |
| | | *Paraplagusia guttata* | Duarte-Ribeiro et al. (CSIROGT1227) | Genome |
| | | *Paraplagusia japonica* | Duarte-Ribeiro et al. (CSIROGT6892) | Genome |
| | | *Paraplagusia japonica* | Duarte-Ribeiro et al. (CSIROHK17) | Genome |
| | Symphurus | | | |
| | | *Symphurus arawak* | Duarte-Ribeiro et al. (USNMAC5ZA85) | Genome |
| | | *Symphurus atricauda* | Duarte-Ribeiro et al. (UNSMT9212) | Genome |
| | | *Symphurus civitatium* | Duarte-Ribeiro et al. (KU5106) | Genome |
| | | *Symphurus diomedeanus* | Duarte-Ribeiro et al. (KU5147) | Genome |
| | | *Symphurus jenynsi* | Duarte-Ribeiro et al. (UNMDP62) | Genome |
| | | *Symphurus microrhynchus* | Duarte-Ribeiro et al. (CSIROIN00579) | Genome |
| | | *Symphurus ommaspilus* | Duarte-Ribeiro et al. (USNMAC2WJ51) | Genome |
| | | *Symphurus plagiusa* | Duarte-Ribeiro et al. (KU1061) | Genome |
| | | Symphurus sp1 | Duarte-Ribeiro et al. (CSIROGT694) | Genome |
| | | Symphurus sp4 | Duarte-Ribeiro et al. (CSIROGT701) | Genome |
| | | Symphurus sp5 | Duarte-Ribeiro et al. (CSIROGT698) | Genome |
| Echeneidae | | | | |
| | Echeneis | | | |
| | | *Echeneis naucrates* | Duarte-Ribeiro et al. (UMSNH34530) | Genome |
| | Remora | | | |
| | | *Remora albescens* | Duarte-Ribeiro et al. (CSIROIN02009) | Genome |
| | | *Remora brachyptera* | Duarte-Ribeiro et al. (CSIROGT8300) | Genome |
| | | *Remora osteochir* | Duarte-Ribeiro et al. (ODU5334) | Genome |
| | | *Remora remora* | Duarte-Ribeiro et al. (USNMAD9NC35) | Genome |
| Holocentridae | | | | |
| | Plectrypops | | | |
| | | *Plectrypops retrospinis* | Duarte-Ribeiro et al. (EPLATE_06_F12) | Genome |
| Istiophoridae | | | | |
| | Istiophorus | | | |
| | | *Istiophorus platypterus* | Duarte-Ribeiro et al. (KU5428) | Genome |
| | Kajikia | | | |
| | | *Kajikia audax* | Duarte-Ribeiro et al. (SIO11345) | Genome |
| | Makaira | | | |
| | | *Makaira nigricans* | Duarte-Ribeiro et al. (KU5430) | Genome |
| | | *Makaira nigricans* | Duarte-Ribeiro et al. (UMSNH25970) | Genome |
| | Tetrapturus | | | |
| | | *Tetrapturus angustirostris* | Duarte-Ribeiro et al. (CSIRO1Tang) | Genome |
| Lactariidae | | | | |
| | Lactarius | | | |
| | | *Lactarius lactarius* | Duarte-Ribeiro et al. (ODU5463) | Genome |
| Leptobramidae | | | | |
| | Leptobrama | | | |
| | | *Leptobrama muelleri* | Duarte-Ribeiro et al. (CSIRO1Lmul) | Genome |
| Lutjanidae | | | | |
| | Gymnocaesio | | | |
| | | *Gymnocaesio gymnoptera* | Duarte-Ribeiro et al. (USNMAB6QR50) | Genome |
| MIS | | | | |
| | Echeneidae | | | |

*Continued on next page*

Table S1 – *Continued from previous page*

| Family | Genera | Species | Accession | Data type |
|---|---|---|---|---|
| | | *Echeneidae Echeneis* | Duarte-Ribeiro et al. (sp1_ODU1036) | Genome |
| | Paralichthyidae | | | |
| | | *Paralichthyidae Paralichthys* | Duarte-Ribeiro et al. (sp3_UMSNH31070) | Genome |
| Menidae | | | | |
| | Mene | | | |
| | | *Mene maculata* | Duarte-Ribeiro et al. (FMNH126719) | Genome |
| Nematistiidae | | | | |
| | Nematistius | | | |
| | | *Nematistius pectoralis* | Duarte-Ribeiro et al. (STRINZOG0226) | Genome |
| Paralichthyidae | | | | |
| | Ancylopsetta | | | |
| | | *Ancylopsetta dendritica* | Duarte-Ribeiro et al. (STRIBFT00263) | Genome |
| | | *Ancylopsetta dilecta* | Duarte-Ribeiro et al. (KU5129) | Genome |
| | | *Ancylopsetta ommata* | Duarte-Ribeiro et al. (KU10) | Genome |
| | | *Ancylopsetta ommata* | Duarte-Ribeiro et al. (KU5223) | Genome |
| | Etropus | | | |
| | | *Etropus cyclosquamus* | Duarte-Ribeiro et al. (KU5241) | Genome |
| | Hippoglossina | | | |
| | | *Hippoglossina stomata* | Duarte-Ribeiro et al. (UW153477) | Genome |
| | | *Hippoglossina tetrophthalma* | Duarte-Ribeiro et al. (USNMAG7PF74) | Genome |
| | Paralichthys | | | |
| | | *Paralichthys albigutta* | Duarte-Ribeiro et al. (USNMAC2WN64) | Genome |
| | | *Paralichthys californicus* | Duarte-Ribeiro et al. (USNMT9175) | Genome |
| | | *Paralichthys dentatus* | Duarte-Ribeiro et al. (KU1446) | Genome |
| | | *Paralichthys lethostigma* | Duarte-Ribeiro et al. (KU5207) | Genome |
| | | *Paralichthys oblongus* | Duarte-Ribeiro et al. (KU1493) | Genome |
| | | *Paralichthys olivaceus* | Duarte-Ribeiro et al. (USNMT10334) | Genome |
| | | *Paralichthys orbignyanus* | Duarte-Ribeiro et al. (UNMDP1506) | Genome |
| | | *Paralichthys squamilentus* | Duarte-Ribeiro et al. (KU5205) | Genome |
| | | *Paralichthys woolmani* | Duarte-Ribeiro et al. (UMSNH11556) | Genome |
| | | *Paralichthys woolmani* | Duarte-Ribeiro et al. (UPRFL1524) | Genome |
| | Pseudorhombus | | | |
| | | *Pseudorhombus argus* | Duarte-Ribeiro et al. (CSIRO1Parg) | Genome |
| | | *Pseudorhombus cfelevatus* | Duarte-Ribeiro et al. (CSIROIN02784) | Genome |
| | | *Pseudorhombus cinnamoneus* | Duarte-Ribeiro et al. (USNMAB6QS40) | Genome |
| | | *Pseudorhombus diplospilus* | Duarte-Ribeiro et al. (CSIROGT4679) | Genome |
| | | *Pseudorhombus diplospilus* | Duarte-Ribeiro et al. (CSIROIN01989) | Genome |
| | | *Pseudorhombus javanicus* | Duarte-Ribeiro et al. (CSIROIN02743) | Genome |
| | | *Pseudorhombus jenynsii* | Duarte-Ribeiro et al. (CSIROGT4425) | Genome |
| | | *Pseudorhombus jenynsii* | Duarte-Ribeiro et al. (CSIROGT5941) | Genome |
| | | *Pseudorhombus pentophthalmus* | Duarte-Ribeiro et al. (KU2481) | Genome |
| | | *Pseudorhombus quinquocellatus* | Duarte-Ribeiro et al. (CSIROGT656) | Genome |
| | | *Pseudorhombus spinosus* | Duarte-Ribeiro et al. (CSIRO1Pspi) | Genome |
| | | *Pseudorhombus tenuirastrum* | Duarte-Ribeiro et al. (CSIROGT3664) | Genome |
| | Tarphops | | | |
| | | *Tarphops oligolepis* | Duarte-Ribeiro et al. (KU2498) | Genome |
| | Thysanopsetta | | | |
| | | *Thysanopsetta naresi* | Duarte-Ribeiro et al. (UNMDP2122) | Genome |

*Continued on next page*

Table S1 – *Continued from previous page*

| Family | Genera | Species | Accession | Data type |
|---|---|---|---|---|
| | Xystreurys | | | |
| | | *Xystreurys liolepis* | Duarte-Ribeiro et al. (USNMT9179) | Genome |
| Pleuronectidae | | | | |
| | Acanthopsetta | | | |
| | | *Acanthopsetta nadeshnyi* | Duarte-Ribeiro et al. (UW118829) | Genome |
| | Atheresthes | | | |
| | | *Atheresthes evermanni* | Duarte-Ribeiro et al. (UW119477) | Genome |
| | | *Atheresthes stomias* | Duarte-Ribeiro et al. (UW151456) | Genome |
| | Cleisthenes | | | |
| | | *Cleisthenes pinetoruxm* | Duarte-Ribeiro et al. (UW118099) | Genome |
| | Clidoderma | | | |
| | | *Clidoderma asperrimum* | Duarte-Ribeiro et al. (UW153538) | Genome |
| | Embassichthys | | | |
| | | *Embassichthys bathybius* | Duarte-Ribeiro et al. (KU2269) | Genome |
| | Eopsetta | | | |
| | | *Eopsetta jordani* | Duarte-Ribeiro et al. (KU555) | Genome |
| | Glyptocephalus | | | |
| | | *Glyptocephalus cynoglossus* | Duarte-Ribeiro et al. (KU1474) | Genome |
| | | *Glyptocephalus zachirus* | Duarte-Ribeiro et al. (USNMT9379) | Genome |
| | Hippoglossoides | | | |
| | | *Hippoglossoides elassodon* | Duarte-Ribeiro et al. (USNMT2090) | Genome |
| | | *Hippoglossoides platessoides* | Duarte-Ribeiro et al. (KU368) | Genome |
| | Hippoglossus | | | |
| | | *Hippoglossus hippoglossus* | Duarte-Ribeiro et al. (KU5417) | Genome |
| | | *Hippoglossus stenolepis* | Duarte-Ribeiro et al. (USNMT3103) | Genome |
| | Hypsopsetta | | | |
| | | *Hypsopsetta guttulata* | Duarte-Ribeiro et al. (USNMT9192) | Genome |
| | Isopsetta | | | |
| | | *Isopsetta isolepis* | Duarte-Ribeiro et al. (KU431) | Genome |
| | Kajikia | | | |
| | | *Kajikia albida* | Duarte-Ribeiro et al. (CSIROGT1414) | Genome |
| | Lepidopsetta | | | |
| | | *Lepidopsetta bilineata* | Duarte-Ribeiro et al. (KU3230) | Genome |
| | | *Lepidopsetta polyxystra* | Duarte-Ribeiro et al. (UW111511) | Genome |
| | | *Lepidopsetta polyxystra* | Duarte-Ribeiro et al. (UW150842) | Genome |
| | Limanda | | | |
| | | *Limanda aspera* | Duarte-Ribeiro et al. (KU385) | Genome |
| | | *Limanda limanda* | Duarte-Ribeiro et al. (KU5418) | Genome |
| | | *Limanda sakhalinensis* | Duarte-Ribeiro et al. (UW152387) | Genome |
| | Liopsetta | | | |
| | | *Liopsetta pinnifasciata* | Duarte-Ribeiro et al. (UW44943) | Genome |
| | Lyopsetta | | | |
| | | *Lyopsetta exilis* | Duarte-Ribeiro et al. (KU3176) | Genome |
| | Microstomus | | | |
| | | *Microstomus kitt* | Duarte-Ribeiro et al. (USNMT5415) | Genome |
| | | *Microstomus pacificus* | Duarte-Ribeiro et al. (KU3209) | Genome |
| | Myzopsetta | | | |
| | | *Myzopsetta ferruginea* | Duarte-Ribeiro et al. (KU1028) | Genome |
| | Oncopterus | | | |
| | | *Oncopterus darwinii* | Duarte-Ribeiro et al. (UNMDP2063) | Genome |

*Continued on next page*

Table S1 – *Continued from previous page*

| Family | Genera | Species | Accession | Data type |
|---|---|---|---|---|
| | Parophrys | | | |
| | | *Parophrys vetulus* | Duarte-Ribeiro et al. (USNMT9375) | Genome |
| | | *Parophrys vetulus* | Duarte-Ribeiro et al. (UW155966) | Genome |
| | Platichthys | | | |
| | | *Platichthys stellatus* | Duarte-Ribeiro et al. (KU428) | Genome |
| | | *Platichthys stellatus* | Duarte-Ribeiro et al. (UW156364) | Genome |
| | Pleuronectes | | | |
| | | *Pleuronectes platessa* | Duarte-Ribeiro et al. (KU1845) | Genome |
| | | *Pleuronectes quadritubercula-tus* | Duarte-Ribeiro et al. (KU381) | Genome |
| | Pleuronichthys | | | |
| | | *Pleuronichthys coenosus* | Duarte-Ribeiro et al. (KU9164) | Genome |
| | | *Pleuronichthys decurrens* | Duarte-Ribeiro et al. (UW116910) | Genome |
| | | *Pleuronichthys ritteri* | Duarte-Ribeiro et al. (USNMT9190) | Genome |
| | Psettichthys | | | |
| | | *Psettichthys melanostictus* | Duarte-Ribeiro et al. (USNMT9373) | Genome |
| | Pseudopleuronectes | | | |
| | | *Pseudopleuronectes americanus* | Duarte-Ribeiro et al. (KU365) | Genome |
| | | *Pseudopleuronectes yokohamae* | Duarte-Ribeiro et al. (USNMT10335) | Genome |
| | Reinhardtius | | | |
| | | *Reinhardtius hippoglossoides* | Duarte-Ribeiro et al. (KU3540) | Genome |
| | Verasper | | | |
| | | *Verasper variegatus* | Duarte-Ribeiro et al. (UW117963) | Genome |
| Poecilopsettidae | | | | |
| | Poecilopsetta | | | |
| | | *Poecilopsetta cfplinthus* | Duarte-Ribeiro et al. (CSIROGT679) | Genome |
| | | *Poecilopsetta cfpraelonga* | Duarte-Ribeiro et al. (CSIROGT1414) | Genome |
| | | *Poecilopsetta natalensis* | Duarte-Ribeiro et al. (CSIROGT5894) | Genome |
| | | *Poecilopsetta plinthus* | Duarte-Ribeiro et al. (KU2472) | Genome |
| | | Poecilopsetta sp1 | Duarte-Ribeiro et al. (CSIROGT651) | Genome |
| Polynemidae | | | | |
| | Filimanus | | | |
| | | *Filimanus perplexa* | Duarte-Ribeiro et al. (CSIROIN02211) | Genome |
| | | *Filimanus xanthonema* | Duarte-Ribeiro et al. (CSIROIN01888) | Genome |
| | Galeoides | | | |
| | | *Galeoides decadactylus* | Duarte-Ribeiro et al. (USNMAD9NF35) | Genome |
| | Pentanemus | | | |
| | | *Pentanemus quinquarius* | Duarte-Ribeiro et al. (STRIBFT05902) | Genome |
| | Polydactylus | | | |
| | | *Polydactylus approximans* | Duarte-Ribeiro et al. (UMSNH10905) | Genome |
| | | *Polydactylus macrochir* | Duarte-Ribeiro et al. (ODU1065) | Genome |
| | | *Polydactylus microstoma* | Duarte-Ribeiro et al. (CSIROIN01802) | Genome |
| | | *Polydactylus multiradiatus* | Duarte-Ribeiro et al. (CSIROGT4464) | Genome |
| | | *Polydactylus octonemus* | Duarte-Ribeiro et al. (USNMT5105) | Genome |
| | | *Polydactylus oligodon* | Duarte-Ribeiro et al. (FL958) | Genome |
| | | *Polydactylus opercularis* | Duarte-Ribeiro et al. (UMSNH23237) | Genome |
| | | *Polydactylus quadrifilis* | Duarte-Ribeiro et al. (ODU1108) | Genome |
| | | *Polydactylus sexfilis* | Duarte-Ribeiro et al. (USNMT6828) | Genome |
| | | *Polydactylus sextarius* | Duarte-Ribeiro et al. (ODU2310) | Genome |
| | | *Polydactylus virginicus* | Duarte-Ribeiro et al. (UMSNH31809) | Genome |

*Continued on next page*

Table S1 – *Continued from previous page*

| Family | Genera | Species | Accession | Data type |
|---|---|---|---|---|
| | Polynemus | | | |
| | | *Polynemus paradiseus* | Duarte-Ribeiro et al. (ODU0646) | Genome |
| Psettodidae | | | | |
| | Psettodes | | | |
| | | *Psettodes belcheri* | Duarte-Ribeiro et al. (ODU0792) | Genome |
| | | *Psettodes erumei* | Duarte-Ribeiro et al. (USNMAG9RD56) | Genome |
| Rachycentridae | | | | |
| | Rachycentron | | | |
| | | *Rachycentron canadum* | Duarte-Ribeiro et al. (USNMT3521) | Genome |
| Rhombosoleidae | | | | |
| | Ammotretis | | | |
| | | *Ammotretis lituratus* | Duarte-Ribeiro et al. (CSIROGT7353) | Genome |
| | | *Ammotretis rostratus* | Duarte-Ribeiro et al. (CSIRO4Aros) | Genome |
| | Colistium | | | |
| | | *Colistium guntheri* | Duarte-Ribeiro et al. (CSIRO1Cgun) | Genome |
| | Mancopsetta | | | |
| | | *Mancopsetta maculata* | Duarte-Ribeiro et al. (CSIROMI393) | Genome |
| | Neoachiropsetta | | | |
| | | *Neoachiropsetta milfordi* | Duarte-Ribeiro et al. (CSIRO2Nmil) | Genome |
| | Rhombosolea | | | |
| | | *Rhombosolea leporina* | Duarte-Ribeiro et al. (CSIRO1vRlep) | Genome |
| | | *Rhombosolea plebeia* | Duarte-Ribeiro et al. (CSIRO1vRple) | Genome |
| | | *Rhombosolea tapirina* | Duarte-Ribeiro et al. (CSIRO6vRtap) | Genome |
| Samaridae | | | | |
| | Plagiopsetta | | | |
| | | *Plagiopsetta glossa* | Duarte-Ribeiro et al. (KU2474) | Genome |
| | Samariscus | | | |
| | | *Samariscus japonicus* | Duarte-Ribeiro et al. (KU2469) | Genome |
| | | *Samariscus triocellatus* | Duarte-Ribeiro et al. (USNMAG5NP35) | Genome |
| | | *Samariscus xenicus* | Duarte-Ribeiro et al. (USNMT2483) | Genome |
| Scombridae | | | | |
| | Rastrelliger | | | |
| | | *Rastrelliger brachysoma* | Duarte-Ribeiro et al. (USNMAG9RJ88) | Genome |
| Scophthalmidae | | | | |
| | Lepidorhombus | | | |
| | | *Lepidorhombus boscii* | Duarte-Ribeiro et al. (KU3496) | Genome |
| | Scophthalmus | | | |
| | | *Scophthalmus aquosus* | Duarte-Ribeiro et al. (KU1252) | Genome |
| Serranidae | | | | |
| | Hypoplectrus | | | |
| | | *Hypoplectrus nigricans* | Duarte-Ribeiro et al. (UPRFL0324) | Genome |
| Soleidae | | | | |
| | Aesopia | | | |
| | | *Aesopia cornuta* | Duarte-Ribeiro et al. (CSIROGT1218) | Genome |
| | Aseraggodes | | | |
| | | *Aseraggodes heemstrai* | Duarte-Ribeiro et al. (KU4997) | Genome |
| | | *Aseraggodes kobensis* | Duarte-Ribeiro et al. (KU2476) | Genome |
| | | *Aseraggodes melanostictus* | Duarte-Ribeiro et al. (CSIROGT4398) | Genome |
| | | *Aseraggodes melanostictus* | Duarte-Ribeiro et al. (KU5719) | Genome |
| | | Aseraggodes sp1 | Duarte-Ribeiro et al. (CSIROGT7521) | Genome |

*Continued on next page*

Table S1 – *Continued from previous page*

| Family | Genera | Species | Accession | Data type |
|---|---|---|---|---|
| | Dexillus | | | |
| | | *Dexillus muelleri* | Duarte-Ribeiro et al. (USNMAC1VI75) | Genome |
| | Heteromycteris | | | |
| | | *Heteromycteris japonicus* | Duarte-Ribeiro et al. (KU2492) | Genome |
| | Monochirus | | | |
| | | *Monochirus hispidus* | Duarte-Ribeiro et al. (ODU0757) | Genome |
| | Pardachirus | | | |
| | | *Pardachirus marmoratus* | Duarte-Ribeiro et al. (FMNHRS1301T06) | Genome |
| | | Pardachirus sp1 | Duarte-Ribeiro et al. (CSIROIN01791) | Genome |
| | Pegusa | | | |
| | | *Pegusa lascaris* | Duarte-Ribeiro et al. (USNMAD9NF57) | Genome |
| | Pseudaesopia | | | |
| | | *Pseudaesopia japonica* | Duarte-Ribeiro et al. (KU2504) | Genome |
| | Solea | | | |
| | | *Solea solea* | Duarte-Ribeiro et al. (KU1846) | Genome |
| | Soleichthys | | | |
| | | *Soleichthys maculosus* | Duarte-Ribeiro et al. (CSIROGT1224) | Genome |
| | | *Soleichthys microcephalus* | Duarte-Ribeiro et al. (CSIROGT1214) | Genome |
| | | *Soleichthys oculofasciatus* | Duarte-Ribeiro et al. (CSIROGT1221) | Genome |
| | | *Soleichthys serpenpellis* | Duarte-Ribeiro et al. (CSIROGT1216) | Genome |
| | Synclidopus | | | |
| | | *Synclidopus macleayanus* | Duarte-Ribeiro et al. (CSIROGT3682) | Genome |
| | Zebrias | | | |
| | | *Zebrias altipinnis* | Duarte-Ribeiro et al. (CSIROIN02961) | Genome |
| | | *Zebrias craticula* | Duarte-Ribeiro et al. (CSIROGT1245) | Genome |
| | | *Zebrias quagga* | Duarte-Ribeiro et al. (CSIROGT1238) | Genome |
| | | *Zebrias scalaris* | Duarte-Ribeiro et al. (CSIROGT3651) | Genome |
| | | *Zebrias zebra* | Duarte-Ribeiro et al. (CSIROHK04) | Genome |
| Sphyraenidae | | | | |
| | Inegocia | | | |
| | | *Inegocia japonica* | Duarte-Ribeiro et al. (USNMT10332) | Genome |
| | Sphyraena | | | |
| | | *Sphyraena argentea* | Duarte-Ribeiro et al. (UMSNH14629) | Genome |
| | | *Sphyraena barracuda* | Duarte-Ribeiro et al. (FL287) | Genome |
| | | *Sphyraena barracuda* | Duarte-Ribeiro et al. (FMNH122802) | Genome |
| | | *Sphyraena ensis* | Duarte-Ribeiro et al. (STRIBFT10051) | Genome |
| | | *Sphyraena guachancho* | Duarte-Ribeiro et al. (UMSNH16904) | Genome |
| | | *Sphyraena idiastes* | Duarte-Ribeiro et al. (UMSNH18299) | Genome |
| | | *Sphyraena jello* | Duarte-Ribeiro et al. (FMNH118454) | Genome |
| | | *Sphyraena picudilla* | Duarte-Ribeiro et al. (USNMAC2WV46) | Genome |
| | | *Sphyraena putnamae* | Duarte-Ribeiro et al. (KU6783) | Genome |
| | | *Sphyraena qenie* | Duarte-Ribeiro et al. (USNMAG9RD64) | Genome |
| | | *Sphyraena viridensis* | Duarte-Ribeiro et al. (USNMAD9NG43) | Genome |
| | | *Sphyraena waitii* | Duarte-Ribeiro et al. (CSIROGT6548) | Genome |
| Synbranchidae | | | | |
| | Synbranchus | | | |
| | | *Synbranchus marmoratus* | Duarte-Ribeiro et al. (EPLATE_23_E09) | Genome |
| Toxotidae | | | | |
| | Toxotes | | | |

*Continued on next page*

Table S1 – *Continued from previous page*

| Family | Genera | Species | Accession | Data type |
|---|---|---|---|---|
| | | *Toxotes chatareus* | Duarte-Ribeiro et al. (ODU1856) | Genome |
| | | *Toxotes jaculatrix* | Duarte-Ribeiro et al. (FMNH119297) | Genome |
| | | *Toxotes lorentzi* | Duarte-Ribeiro et al. (ODU1858) | Genome |
| Xiphiidae | | | | |
| | Xiphias | | | |
| | | *Xiphias gladius* | Duarte-Ribeiro et al. (SRR3213608) | Genome |
| Zenarchopteridae | | | | |
| | Zenarchopterus | | | |
| | | *Zenarchopterus dispar* | Duarte-Ribeiro et al. (EPLATE_31_A11) | Genome |

Table S2: List of species examined of the Protacanthopterygii dataset

| Order | Family | Genera | Species | Accession | Data type |
|---|---|---|---|---|---|
| Alepocephaliformes | Platytroctidae | | | | |
| | | Holtbyrnia | | | |
| | | | *Holtbyrnia laticauda* | This study (EPLATE_28_H06) | Genome |
| | | Normichthys | | | |
| | | | *Normichthys yahganorum* | This study (EPLATE_28_H07) | Genome |
| | | Persparsia | | | |
| | | | *Persparsia kopua* | This study (EPLATE_46_D03) | Genome |
| Argentiniformes | | | | | |
| | Argentinidae | | | | |
| | | Argentina | | | |
| | | | *Argentina australiae* | This study (EPLATE_46_F03) | Genome |
| | | | *Argentina silus* | SRR11679483 | Genome |
| | | | Argentina sp | SRR5997680 | Transcriptome |
| | | | Argentina sp | SRX3153196 | Transcriptome |
| | | | *Argentina sphyraena* | SRR11537143 | Genome |
| | | Glossanodon | | | |
| | | | Glossanodon spW1 | This study (EPLATE_46_E03) | Genome |
| | Bathylagidae | | | | |
| | | Bathylagichthys | | | |
| | | | *Bathylagichthys australis* | This study (EPLATE_46_H03) | Genome |
| | | | *Bathylagichthys longipinnis* | This study (EPLATE_46_G03) | Genome |
| | | | *Bathylagichthys problematicus* | This study (EPLATE_46_A04) | Genome |
| | | Melanolagus | | | |
| | | | *Melanolagus bericoides* | This study (EPLATE_46_C04) | Genome |
| | | | Melanolagus sp2 | This study (EPLATE_46_B04) | Genome |

*Continued on next page*

Table S2 – *Continued from previous page*

| Order | Family | Genera | Species | Accession | Data type |
|---|---|---|---|---|---|
| | Microstomatidae | | | | |
| | | Microstoma | | | |
| | | | *Microstoma microstoma* | This study (EPLATE_46_D04) | Genome |
| | | Nansenia | | | |
| | | | *Nansenia antarctica* | This study (EPLATE_46_E04) | Genome |
| | Opisthoproctidae | | | | |
| | | Monacoa | | | |
| | | | Monacoa sp1 | This study (EPLATE_46_G04) | Genome |
| | | Opisthoproctus | | | |
| | | | *Opisthoproctus soleatus* | ERR3332507 | Genome |
| | | Winteria | | | |
| | | | *Winteria telescopa* | This study (EPLATE_46_H04) | Genome |
| Ateleopodiformes | | | | | |
| | Ateleopodidae | | | | |
| | | Ateleopus | | | |
| | | | *Ateleopus japonicus* | This study (EPLATE_26_E02) | Genome |
| | | Ijimaia | | | |
| | | | *Ijimaia dofleini* | This study (EPLATE_26_E03) | Genome |
| | | | *Ijimaia dofleini* | This study (EPLATE_46_A05) | Genome |
| | | | Ijimaia sp1 | This study (EPLATE_46_C05) | Genome |
| Aulopiformes | | | | | |
| | Aulopidae | | | | |
| | | Hime | | | |
| | | | *Hime caudizoma* | This study (EPLATE_60_B05) | Genome |
| | | | *Hime curtirostris* | This study (EPLATE_61_B03) | Genome |
| | Bathysauroididae | | | | |
| | | Bathysauroides | | | |
| | | | *Bathysauroides gigas* | This study (EPLATE_46_E05) | Genome |
| | Paralepididae | | | | |
| | | Lestidium | | | |
| | | | *Lestidium atlanticum* | This study (EPLATE_61_C05) | Genome |
| Clupeiformes | | | | | |
| | Clupeidae | | | | |
| | | Alosa | | | |
| | | | *Alosa alosa* | PRJNA256955 | Transcriptome |
| | | Amblygaster | | | |
| | | | *Amblygaster clupeoides* | SRX3153191 | Transcriptome |

*Continued on next page*

Table S2 – *Continued from previous page*

| Order | Family | Genera | Species | Accession | Data type |
|---|---|---|---|---|---|
| | | Clupea | | | |
| | | | *Clupea harengus* | GCA_000966335.1 | Genome |
| | Engraulidae | | | | |
| | | Coilia | | | |
| | | | *Coilia nasus* | SRX3153210 | Transcriptome |
| | | Engraulis | | | |
| | | | *Engraulis encrasicolus* | PRJNA261165 | Transcriptome |
| Esociformes | | | | | |
| | Esocidae | | | | |
| | | Esox | | | |
| | | | *Esox lucius* | GCA_000721915.2 | Genome |
| | | | *Esox reichertii* | This study (EPLATE_44_C07) | Genome |
| | Umbridae | | | | |
| | | Umbra | | | |
| | | | *Umbra pygmaea* | PRJNA257000 | Transcriptome |
| Galaxiiformes | | | | | |
| | Galaxiidae | | | | |
| | | Galaxias | | | |
| | | | *Galaxias maculatus* | Hughes et al. (2018) | Genome |
| | | | *Galaxias truttaceus* | This study (EPLATE_25_B09) | Genome |
| | | Galaxiella | | | |
| | | | *Galaxiella nigrostriata* | SRX3153920 | Transcriptome |
| | | Lovettia | | | |
| | | | *Lovettia sealii* | This study (EPLATE_25_B10) | Genome |
| Lepidogalaxiiformes | | | | | |
| | Lepidogalaxiidae | | | | |
| | | Lepidogalaxias | | | |
| | | | *Lepidogalaxias salamandroides* | SRX3153919 | Transcriptome |
| Myctophiformes | | | | | |
| | Myctophidae | | | | |
| | | Benthosema | | | |
| | | | *Benthosema fibulatum* | This study (EPLATE_49_A06) | Genome |
| | | Diaphus | | | |
| | | | *Diaphus hudsoni* | This study (EPLATE_49_G02) | Genome |
| | | | *Diaphus termophilus* | This study (EPLATE_49_E01) | Genome |
| | | | *Diaphus whitleyi* | This study (EPLATE_49_G09) | Genome |
| Osmeriformes | | | | | |
| | Osmeridae | | | | |
| | | Mallotus | | | |
| | | | *Mallotus villosus* | GCA_903064625.1 | Genome |
| | | Osmerus | | | |

*Continued on next page*

Table S2 – *Continued from previous page*

| Order | Family | Genera | Species | Accession | Data type |
|---|---|---|---|---|---|
| | | | *Osmerus eperlanus* | GCA_900302275.1 | Genome |
| | | | *Osmerus mordax* | This study (EPLATE_44_E11) | Genome |
| | | Spirinchus | | | |
| | | | *Spirinchus thaleichthys* | SRR2758850 | Transcriptome |
| | | Thaleichthys | | | |
| | | | *Thaleichthys pacificus* | SRR1786619 | Genome |
| | Plecoglossidae | | | | |
| | | Plecoglossus | | | |
| | | | *Plecoglossus altivelis* | This study (EPLATE_44_G03) | Genome |
| | Salangidae | | | | |
| | | Protosalanx | | | |
| | | | *Protosalanx hyalocranius* | PRJNA328051 | Genome |
| | | Salanx | | | |
| | | | *Salanx chinensis* | GCA_010882115.1 | Genome |
| | | | Salanx sp1 | This study (EPLATE_64_D11) | Genome |
| Salmoniformes | | | | | |
| | Salmonidae | | | | |
| | | Brachymystax | | | |
| | | | *Brachymystax lenok* | SRR8873459 | Transcriptome |
| | | Coregonus | | | |
| | | | Coregonus 22 | | |
| | | | *Coregonus artedi* | SRR9718093 | Genome |
| | | | *Coregonus clupeaformis* | PRJNA256999 | Transcriptome |
| | | | *Coregonus lavaretus* | SRR9951409 | Genome |
| | | | *Coregonus palaea* | SRR2175595 | Transcriptome |
| | | | Coregonus sp | GCA_902810595.1 | Genome |
| | | Hucho | | | |
| | | | *Hucho hucho* | GCA_003317085.1 | Genome |
| | | | *Hucho taimen* | ERR1857179 | Transcriptome |
| | | Oncorhynchus | | | |
| | | | *Oncorhynchus keta* | GCF_012931545.1 | Genome |
| | | | *Oncorhynchus kisutch* | GCF_002021735.2 | Genome |
| | | | *Oncorhynchus masou* | DRR239371 | Transcriptome |
| | | | *Oncorhynchus mykiss* | PRJEB4421 | Genome |
| | | | *Oncorhynchus nerka* | GCF_006149115.1 | Genome |
| | | | *Oncorhynchus tshawytscha* | This study (EPLATE_62_C09) | Genome |
| | | Salmo | | | |
| | | | *Salmo ischchan* | SRR6854514 | Genome |
| | | | *Salmo marmoratus* | SRR7687456 | Transcriptome |
| | | | *Salmo salar* | GCA_000233375.4 | Genome |

*Continued on next page*

Table S2 – *Continued from previous page*

| Order | Family | Genera | Species | Accession | Data type |
|---|---|---|---|---|---|
| | | | *Salmo trutta* | GCF_901001165.1 | Genome |
| | | Salvelinus | | | |
| | | | *Salvelinus alpinus* | SRR8477003 | Genome |
| | | | *Salvelinus fontinalis* | PRJNA256998 | Transcriptome |
| | | | *Salvelinus namaycush* | SRR11144341 | Genome |
| | | | Salvelinus sp | GCF_002910315.2 | Genome |
| | | Thymallus | | | |
| | | | *Thymallus thymallus* | PRJNA256970 | Transcriptome |
| Stomiatiformes | | | | | |
| | Gonostomatidae | | | | |
| | | Diplophos | | | |
| | | | *Diplophos australis* | This study (EPLATE_53_G09) | Genome |
| | | | *Diplophos rebainsi* | This study (EPLATE_53_F09) | Genome |
| | | | *Diplophos taenia* | This study (EPLATE_18_F09) | Genome |
| | | Margrethia | | | |
| | | | *Margrethia valentinae* | This study (EPLATE_53_D10) | Genome |
| | | Sigmops | | | |
| | | | *Sigmops bathyphilus* | This study (EPLATE_53_C10) | Genome |
| | | | *Sigmops elongatus* | This study (EPLATE_53_H09) | Genome |
| | | | *Sigmops elongatus* | This study (EPLATE_53_A10) | Genome |
| | Phosichthyidae | | | | |
| | | Ichthyococcus | | | |
| | | | *Ichthyococcus australis* | This study (EPLATE_53_E10) | Genome |
| | | Phosichthys | | | |
| | | | *Phosichthys argenteus* | This study (EPLATE_53_A11) | Genome |
| | | Polymetme | | | |
| | | | *Polymetme corythaeola* | This study (EPLATE_53_G10) | Genome |
| | | | *Polymetme illustris* | This study (EPLATE_53_H10) | Genome |
| | | Vinciguerria | | | |
| | | | *Vinciguerria attenuata* | This study (EPLATE_53_B11) | Genome |
| | | | *Vinciguerria attenuata* | This study (EPLATE_53_D11) | Genome |
| | | | *Vinciguerria nimbaria* | This study (EPLATE_53_C11) | Genome |
| | | Woodsia | | | |
| | | | *Woodsia meyerwaardeni* | This study (EPLATE_53_E11) | Genome |

*Continued on next page*

Table S2 – *Continued from previous page*

| Order | Family | Genera | Species | Accession | Data type |
|---|---|---|---|---|---|
| | Sternoptychidae | | | | |
| | | Argyripnus | | | |
| | | | *Argyripnus atlanticus* | This study (EPLATE_18_H06) | Genome |
| | | | *Argyripnus iridescens* | This study (EPLATE_53_F08) | Genome |
| | | Argyropelecus | | | |
| | | | *Argyropelecus aculeatus* | This study (EPLATE_33_F02) | Genome |
| | | | *Argyropelecus affinis* | This study (EPLATE_18_F08) | Genome |
| | | | *Argyropelecus gigas* | This study (EPLATE_53_B08) | Genome |
| | | | *Argyropelecus hemigymnus* | This study (EPLATE_53_D08) | Genome |
| | | | *Argyropelecus olfersii* | This study (EPLATE_18_H09) | Genome |
| | | | *Argyropelecus sladeni* | This study (EPLATE_53_E08) | Genome |
| | | Maurolicus | | | |
| | | | *Maurolicus japonicus* | This study (EPLATE_56_F05) | Genome |
| | | | *Maurolicus mucronatus* | SRR6008916 | Transcriptome |
| | | | *Maurolicus muelleri* | This study (EPLATE_24_C05) | Genome |
| | | Polyipnus | | | |
| | | | *Polyipnus ruggeri* | This study (EPLATE_53_B09) | Genome |
| | | Sternoptyx | | | |
| | | | *Sternoptyx diaphana* | This study (EPLATE_18_E07) | Genome |
| | | | *Sternoptyx pseudodiaphana* | This study (EPLATE_18_G07) | Genome |
| | Stomiidae | | | | |
| | | Astronesthes | | | |
| | | | *Astronesthes boulengeri* | This study (EPLATE_53_F11) | Genome |
| | | | *Astronesthes illuminatus* | This study (EPLATE_53_G11) | Genome |
| | | Bathophilus | | | |
| | | | *Bathophilus abarbatus* | This study (EPLATE_53_G12) | Genome |
| | | Borostomias | | | |
| | | | *Borostomias antarcticus* | GCA_900323325.1 | Genome |
| | | Chauliodus | | | |
| | | | *Chauliodus sloani* | This study (EPLATE_18_A09) | Genome |
| | | Malacosteus | | | |

*Continued on next page*

Table S2 – *Continued from previous page*

| **Order** | **Family** | **Genera** | **Species** | **Accession** | **Data type** |
|---|---|---|---|---|---|
| | | | *Malacosteus australis* | This study (EPLATE_53_C12) | Genome |
| | | | *Malacosteus niger* | This study (EPLATE_18_A10) | Genome |
| | | Neonesthes | | | |
| | | | *Neonesthes microcephalus* | This study (EPLATE_53_B12) | Genome |
| | | Pachystomias | | | |
| | | | *Pachystomias microdon* | SRR5069729 | Transcriptome |
| | | Photostomias | | | |
| | | | *Photostomias tantillux* | This study (EPLATE_18_D09) | Genome |
| | | Rhadinesthes | | | |
| | | | *Rhadinesthes decimus* | This study (EPLATE_53_A12) | Genome |
| | | Stomias | | | |
| | | | *Stomias affinis* | This study (EPLATE_18_G08) | Genome |
| | | | *Stomias boa* | This study (EPLATE_24_B12) | Genome |
| | | | *Stomias gracilis* | This study (EPLATE_53_F12) | Genome |
| Zeiformes | | | | | |
| | Oreosomatidae | | | | |
| | | Allocyttus | | | |
| | | | *Allocyttus verrucosus* | This study (EPLATE_33_E07) | Genome |
| | | Oreosoma | | | |
| | | | *Oreosoma atlanticum* | This study (EPLATE_53_A08) | Genome |
| | | Pseudocyttus | | | |
| | | | *Pseudocyttus maculatus* | This study (EPLATE_53_G07) | Genome |

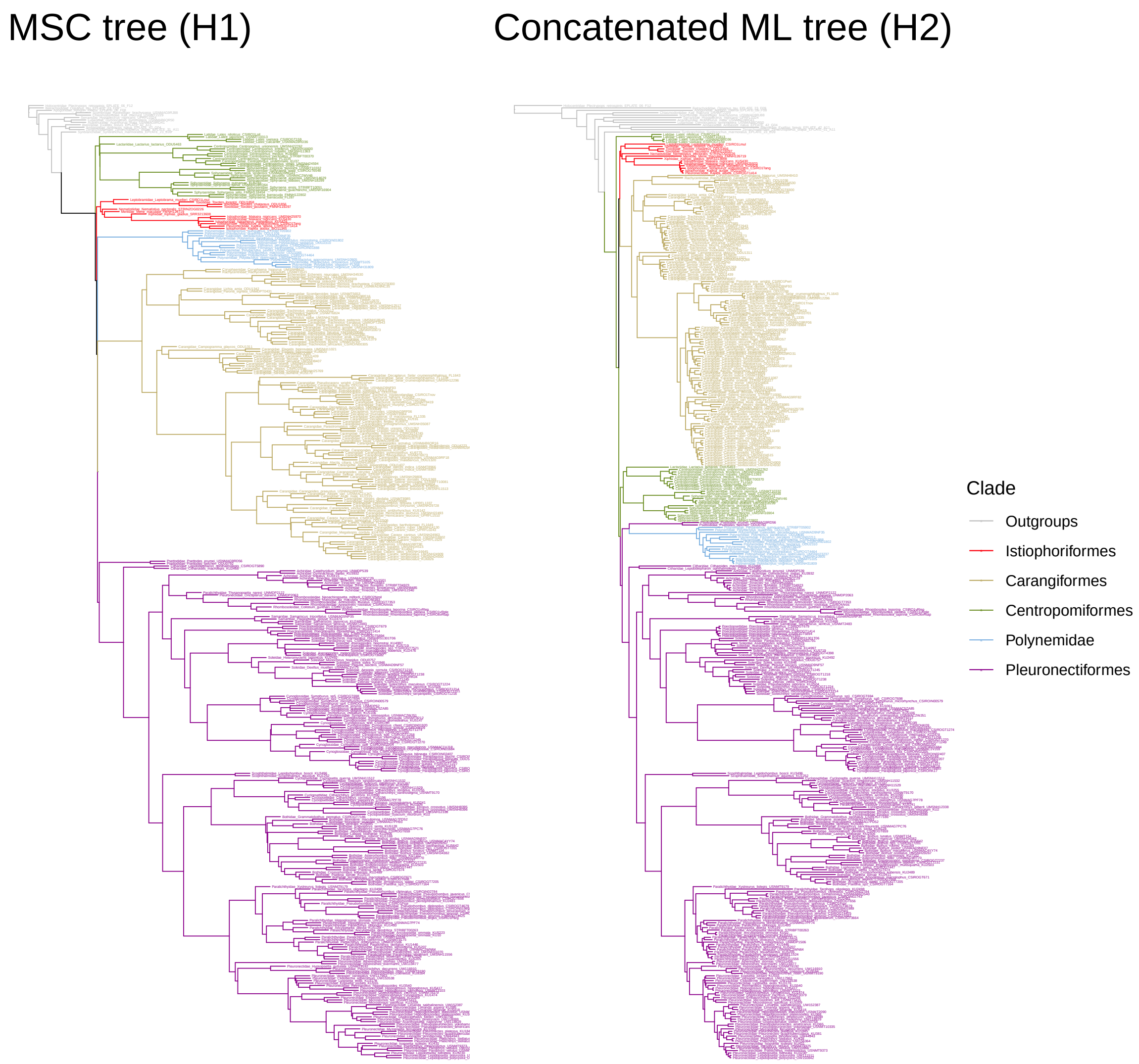

Figure S1: Complete phylogenetic hypotheses for the Carangaria dataset. The left panel shows the MSC tree obtained with ASTRAL (H1) and the right panel shows the concatenated ML tree obtained with IQ-TREE (H2)

Table S3: List of hypotheses tested with GGI for the Protacanthopterygii dataset. Outgroup is composed by the orders Lepidogalaxiiformes, Alepocephaliformes, and Clupeiformes

| **Label** | **Hypothesis** | **Authors** |
|---|---|---|
| H1 | (Outgroup,(Eso_salmo,(Argentiniformes,(Osme_Stomia,(Neoteleostei,Galaxiiformes))))); | Miya and Nishida (2015) |

*Continued on next page*

Table S3 – *Continued from previous page*

| Label | Hypothesis | Authors |
|---|---|---|
| H2 | (Outgroup,((Eso_salmo,Argentiniformes),(Osme_Stomia,(Neoteleostei,Galaxiiformes)))); | Near et al. (2012); Ybazeta (2012); Hughes et al. (2018) |
| H3 | (Outgroup,(Argentiniformes,(Eso_salmo,(Osme_Stomia,(Neoteleostei,Galaxiiformes))))); | |
| H4 | (Outgroup,((Neoteleostei,Galaxiiformes),(Osme_Stomia,(Eso_salmo,Argentiniformes)))); | |
| H5 | (Outgroup,(Galaxiiformes,(Neoteleostei,(Osme_Stomia,(Eso_salmo,Argentiniformes))))); | |
| H6 | (Outgroup,((Eso_salmo,Argentiniformes),(Neoteleostei,(Osme_Stomia,Galaxiiformes)))); | López et al. (2004) |
| H7 | (Outgroup,(Osme_Stomia,((Eso_salmo,Argentiniformes),(Neoteleostei,Galaxiiformes)))); | |
| H8 | (Outgroup,(Eso_salmo,(Osme_Stomia,(Argentiniformes,(Neoteleostei,Galaxiiformes))))); | |
| H9 | (Outgroup,(Eso_salmo,((Neoteleostei,Galaxiiformes),(Osme_Stomia,Argentiniformes)))); | |
| H10 | (Outgroup,(Eso_salmo,(Argentiniformes,(Neoteleostei,(Osme_Stomia,Galaxiiformes))))); | |
| H11 | (Outgroup,(Argentiniformes,(Osme_Stomia,(Eso_salmo,(Neoteleostei,Galaxiiformes))))); | |
| H12 | (Outgroup,((Neoteleostei,Galaxiiformes),(Argentiniformes,(Osme_Stomia,Eso_salmo)))); | |
| H13 | (Outgroup,(Galaxiiformes,(Osme_Stomia,((Eso_salmo,Argentiniformes),Neoteleostei)))); | |
| H14 | (Outgroup,(Neoteleostei,(Galaxiiformes,(Osme_Stomia,(Eso_salmo,Argentiniformes))))); | Li et al. (2010) |
| H15 | (Outgroup,(Osme_Stomia,((Eso_salmo,(Neoteleostei,Galaxiiformes)),Argentiniformes))); | |
| H16 | (Outgroup,(Eso_salmo,((Argentiniformes,Neoteleostei),(Osme_Stomia,Galaxiiformes)))); | |
| H17 | (Outgroup,(Eso_salmo,(Argentiniformes,(Galaxiiformes,(Osme_Stomia,Neoteleostei))))); | |
| H18 | (Outgroup,(Argentiniformes,((Neoteleostei,Galaxiiformes),(Osme_Stomia,Eso_salmo)))); | |
| H19 | (Outgroup,(Galaxiiformes,(Neoteleostei,(Argentiniformes,(Osme_Stomia,Eso_salmo))))); | |
| H20 | (Outgroup,(Osme_Stomia,(Eso_salmo,(Argentiniformes,(Neoteleostei,Galaxiiformes))))); | |
| H21 | (Outgroup,(Eso_salmo,(Osme_Stomia,((Argentiniformes,Galaxiiformes),Neoteleostei)))); | |
| H22 | (Outgroup,(((Eso_salmo,Argentiniformes),Neoteleostei),(Osme_Stomia,Galaxiiformes))); | |
| H23 | (Outgroup,((Neoteleostei,Galaxiiformes),(Eso_salmo,(Osme_Stomia,Argentiniformes)))); | |
| H24 | (Outgroup,(Galaxiiformes,((Eso_salmo,Argentiniformes),(Osme_Stomia,Neoteleostei)))); | Mirande (2017) |
| H25 | (Outgroup,(Neoteleostei,(Osme_Stomia,((Eso_salmo,Galaxiiformes),Argentiniformes)))); | |
| H26 | (Outgroup,(Neoteleostei,(Osme_Stomia,((Eso_salmo,Argentiniformes),Galaxiiformes)))); | |
| H27 | (Outgroup,(Osme_Stomia,(((Eso_salmo,Argentiniformes),Galaxiiformes),Neoteleostei))); | |
| H28 | (Outgroup,(Argentiniformes,((Eso_salmo,Neoteleostei),(Osme_Stomia,Galaxiiformes)))); | |
| H29 | (Outgroup,(Eso_salmo,(Neoteleostei,(Argentiniformes,(Osme_Stomia,Galaxiiformes))))); | Nelson et al. (2016); Hughes et al. (2018) |
| H30 | (Outgroup,(Eso_salmo,(Osme_Stomia,((Argentiniformes,Neoteleostei),Galaxiiformes)))); | |
| H31 | (Outgroup,(Neoteleostei,((Eso_salmo,Argentiniformes),(Osme_Stomia,Galaxiiformes)))); | Campbell et al. (2013) |
| H32 | (Outgroup,(Neoteleostei,(Eso_salmo,(Osme_Stomia,(Argentiniformes,Galaxiiformes))))); | |
| H33 | (Outgroup,((Argentiniformes,(Neoteleostei,Galaxiiformes)),(Osme_Stomia,Eso_salmo))); | |
| H34 | (Outgroup,(Argentiniformes,((Eso_salmo,Galaxiiformes),(Osme_Stomia,Neoteleostei)))); | |
| H35 | (Outgroup,(Eso_salmo,(Neoteleostei,(Osme_Stomia,(Argentiniformes,Galaxiiformes))))); | |
| H36 | (Outgroup,(Neoteleostei,(Eso_salmo,(Argentiniformes,(Osme_Stomia,Galaxiiformes))))); | |
| H37 | (Outgroup,(((Eso_salmo,Argentiniformes),Galaxiiformes),(Osme_Stomia,Neoteleostei))); | |
| H38 | (Outgroup,(((Eso_salmo,Galaxiiformes),Neoteleostei),(Osme_Stomia,Argentiniformes))); | |
| H39 | (Outgroup,((Eso_salmo,(Argentiniformes,Galaxiiformes)),(Osme_Stomia,Neoteleostei))); | |

*Continued on next page*

Table S3 – *Continued from previous page*

| Label | Hypothesis | Authors |
|---|---|---|
| H40 | (Outgroup,((Eso_salmo,Argentiniformes),(Galaxiiformes,(Osme_Stomia,Neoteleostei)))); | |
| H41 | (Outgroup,(Argentiniformes,(Eso_salmo,(Neoteleostei,(Osme_Stomia,Galaxiiformes))))); | |
| H42 | (Outgroup,(Eso_salmo,(Neoteleostei,(Galaxiiformes,(Osme_Stomia,Argentiniformes))))); | |
| H43 | (Outgroup,(Galaxiiformes,(Eso_salmo,(Argentiniformes,(Osme_Stomia,Neoteleostei))))); | |
| H44 | (Outgroup,(Galaxiiformes,(Neoteleostei,(Eso_salmo,(Osme_Stomia,Argentiniformes))))); | |
| H45 | (Outgroup,(Galaxiiformes,(Osme_Stomia,(Eso_salmo,(Argentiniformes,Neoteleostei))))); | |
| H46 | (Outgroup,(Neoteleostei,((Argentiniformes,Galaxiiformes),(Osme_Stomia,Eso_salmo)))); | |
| H47 | (Outgroup,(Neoteleostei,((Eso_salmo,Galaxiiformes),(Osme_Stomia,Argentiniformes)))); | |
| H48 | (Outgroup,((Argentiniformes,Neoteleostei),(Eso_salmo,(Osme_Stomia,Galaxiiformes)))); | |
| H49 | (Outgroup,((Eso_salmo,Galaxiiformes),(Argentiniformes,(Osme_Stomia,Neoteleostei)))); | |
| H50 | (Outgroup,(Argentiniformes,(Osme_Stomia,((Eso_salmo,Neoteleostei),Galaxiiformes)))); | |
| H51 | (Outgroup,(Galaxiiformes,(Argentiniformes,(Osme_Stomia,(Eso_salmo,Neoteleostei))))); | |
| H52 | (Outgroup,(Neoteleostei,(Galaxiiformes,(Argentiniformes,(Osme_Stomia,Eso_salmo))))); | |
| H53 | (Outgroup,(Osme_Stomia,(((Eso_salmo,Galaxiiformes),Argentiniformes),Neoteleostei))); | |
| H54 | (Outgroup,(Osme_Stomia,(((Eso_salmo,Neoteleostei),Galaxiiformes),Argentiniformes))); | |
| H55 | (Outgroup,(((Eso_salmo,Galaxiiformes),Argentiniformes),(Osme_Stomia,Neoteleostei))); | Betancur et al. (2017) |
| H56 | (Outgroup,(((Eso_salmo,Neoteleostei),Argentiniformes),(Osme_Stomia,Galaxiiformes))); | |
| H57 | (Outgroup,((Argentiniformes,Galaxiiformes),(Osme_Stomia,(Eso_salmo,Neoteleostei)))); | |
| H58 | (Outgroup,((Eso_salmo,Galaxiiformes),(Neoteleostei,(Osme_Stomia,Argentiniformes)))); | |
| H59 | (Outgroup,(Argentiniformes,(Neoteleostei,(Eso_salmo,(Osme_Stomia,Galaxiiformes))))); | |
| H60 | (Outgroup,(Argentiniformes,(Neoteleostei,(Osme_Stomia,(Eso_salmo,Galaxiiformes))))); | |
| H61 | (Outgroup,(Eso_salmo,((Argentiniformes,Galaxiiformes),(Osme_Stomia,Neoteleostei)))); | |
| H62 | (Outgroup,(Eso_salmo,(Galaxiiformes,(Osme_Stomia,(Argentiniformes,Neoteleostei))))); | |
| H63 | (Outgroup,(Galaxiiformes,(Argentiniformes,(Eso_salmo,(Osme_Stomia,Neoteleostei))))); | |
| H64 | (Outgroup,(Galaxiiformes,(Eso_salmo,(Osme_Stomia,(Argentiniformes,Neoteleostei))))); | |
| H65 | (Outgroup,(Osme_Stomia,(((Eso_salmo,Argentiniformes),Neoteleostei),Galaxiiformes))); | |
| H66 | (Outgroup,(Osme_Stomia,(((Eso_salmo,Neoteleostei),Argentiniformes),Galaxiiformes))); | |
| H67 | (Outgroup,(((Argentiniformes,Galaxiiformes),Neoteleostei),(Osme_Stomia,Eso_salmo))); | |
| H68 | (Outgroup,((Argentiniformes,Galaxiiformes),(Eso_salmo,(Osme_Stomia,Neoteleostei)))); | |
| H69 | (Outgroup,((Argentiniformes,Neoteleostei),(Osme_Stomia,(Eso_salmo,Galaxiiformes)))); | |
| H70 | (Outgroup,((Eso_salmo,(Argentiniformes,Neoteleostei)),(Osme_Stomia,Galaxiiformes))); | |
| H71 | (Outgroup,((Eso_salmo,Neoteleostei),(Argentiniformes,(Osme_Stomia,Galaxiiformes)))); | |
| H72 | (Outgroup,(Argentiniformes,(Eso_salmo,(Galaxiiformes,(Osme_Stomia,Neoteleostei))))); | |
| H73 | (Outgroup,(Argentiniformes,(Neoteleostei,(Galaxiiformes,(Osme_Stomia,Eso_salmo))))); | |
| H74 | (Outgroup,(Argentiniformes,(Osme_Stomia,((Eso_salmo,Galaxiiformes),Neoteleostei)))); | |
| H75 | (Outgroup,(Eso_salmo,(Galaxiiformes,(Argentiniformes,(Osme_Stomia,Neoteleostei))))); | |
| H76 | (Outgroup,(Eso_salmo,(Galaxiiformes,(Neoteleostei,(Osme_Stomia,Argentiniformes))))); | |
| H77 | (Outgroup,(Galaxiiformes,((Argentiniformes,Neoteleostei),(Osme_Stomia,Eso_salmo)))); | |
| H78 | (Outgroup,(Galaxiiformes,((Eso_salmo,Neoteleostei),(Osme_Stomia,Argentiniformes)))); | |
| H79 | (Outgroup,(Galaxiiformes,(Argentiniformes,(Neoteleostei,(Osme_Stomia,Eso_salmo))))); | |
| H80 | (Outgroup,(Neoteleostei,(Argentiniformes,(Galaxiiformes,(Osme_Stomia,Eso_salmo))))); | |
| H81 | (Outgroup,(Neoteleostei,(Argentiniformes,(Osme_Stomia,(Eso_salmo,Galaxiiformes))))); | |
| H82 | (Outgroup,((Eso_salmo,(Neoteleostei,Galaxiiformes)),(Osme_Stomia,Argentiniformes))); | |
| H83 | (Outgroup,((Eso_salmo,Neoteleostei),(Osme_Stomia,(Argentiniformes,Galaxiiformes)))); | |
| H84 | (Outgroup,(Galaxiiformes,(Osme_Stomia,((Eso_salmo,Neoteleostei),Argentiniformes)))); | |
| H85 | (Outgroup,(Neoteleostei,(Eso_salmo,(Galaxiiformes,(Osme_Stomia,Argentiniformes))))); | |
| H86 | (Outgroup,(Neoteleostei,(Osme_Stomia,(Eso_salmo,(Argentiniformes,Galaxiiformes))))); | |
| H87 | (Outgroup,(Osme_Stomia,(((Eso_salmo,Galaxiiformes),Neoteleostei),Argentiniformes))); | |

*Continued on next page*

Table S3 – *Continued from previous page*

| Label | Hypothesis | Authors |
|---|---|---|
| H88 | (Outgroup,(Osme_Stomia,((Eso_salmo,(Argentiniformes,Galaxiiformes)),Neoteleostei))); | |
| H89 | (Outgroup,(Osme_Stomia,((Eso_salmo,Galaxiiformes),(Argentiniformes,Neoteleostei)))); | |
| H90 | (Outgroup,(Osme_Stomia,((Eso_salmo,Neoteleostei),(Argentiniformes,Galaxiiformes)))); | |
| H91 | (Outgroup,(((Argentiniformes,Neoteleostei),Galaxiiformes),(Osme_Stomia,Eso_salmo))); | |
| H92 | (Outgroup,((Argentiniformes,Neoteleostei),(Galaxiiformes,(Osme_Stomia,Eso_salmo)))); | |
| H93 | (Outgroup,((Eso_salmo,Neoteleostei),(Galaxiiformes,(Osme_Stomia,Argentiniformes)))); | |
| H94 | (Outgroup,(Argentiniformes,(Galaxiiformes,(Neoteleostei,(Osme_Stomia,Eso_salmo))))); | |
| H95 | (Outgroup,(Neoteleostei,(Galaxiiformes,(Eso_salmo,(Osme_Stomia,Argentiniformes))))); | |
| H96 | (Outgroup,(Osme_Stomia,((Eso_salmo,(Argentiniformes,Neoteleostei)),Galaxiiformes))); | |
| H97 | (Outgroup,(Osme_Stomia,(Eso_salmo,((Argentiniformes,Galaxiiformes),Neoteleostei)))); | |
| H98 | (Outgroup,(((Eso_salmo,Neoteleostei),Galaxiiformes),(Osme_Stomia,Argentiniformes))); | |
| H99 | (Outgroup,((Argentiniformes,Galaxiiformes),(Neoteleostei,(Osme_Stomia,Eso_salmo)))); | |
| H100 | (Outgroup,((Eso_salmo,Galaxiiformes),(Osme_Stomia,(Argentiniformes,Neoteleostei)))); | |
| H101 | (Outgroup,(Argentiniformes,(Galaxiiformes,(Eso_salmo,(Osme_Stomia,Neoteleostei))))); | |
| H102 | (Outgroup,(Argentiniformes,(Galaxiiformes,(Osme_Stomia,(Eso_salmo,Neoteleostei))))); | |
| H103 | (Outgroup,(Galaxiiformes,(Eso_salmo,(Neoteleostei,(Osme_Stomia,Argentiniformes))))); | |
| H104 | (Outgroup,(Neoteleostei,(Argentiniformes,(Eso_salmo,(Osme_Stomia,Galaxiiformes))))); | |
| H105 | (Outgroup,(Osme_Stomia,(Eso_salmo,((Argentiniformes,Neoteleostei),Galaxiiformes)))); | |

Table S4: List of features used based on alignment and tree information for training machine learning models. Features were obtained with the "qcutil" command line, a utility of the program "fishlifeqc"

| Variable name | Description |
|---|---|
| nheaders | number of sequences at the alignment |
| pis | percentage of parsimony-informative sites |
| vars | percentage of variable sites |
| seq_len | alignment length |
| seq_len_nogap | percentage of number of sites without gaps |
| gap_prop | proportion of gaps characters for the alignment matrix |
| nogap_prop | proportion of non-gap characters for the alignment matrix |
| gc_mean | mean of GC-content per sequence |
| gc_var | variance of GC-content per sequence |
| gap_mean | mean of gap percentage per sequence |
| gap_var | variance of gap percentage per sequence |
| pi_mean | mean of pairwise identity |
| pi_std | standard deviation of pairwise identity |
| entropy | average of Shannon Entropy (Shannon, 1948) for all columns |
| singletons | a site containing one type of nucleotide ocurring multiple times |
| invariants | percentage of a site containing same nucleotides in all sequences |
| pattern | number of unique site pattern for all columns |
| total_tree_len | sum of all branch lengths in the tree |
| treeness | sum of internal branch lengths devided by 'total_tree_len' (as described by Phillips and Penny (2003)) |
| inter_len_mean | mean of internal branch lengths in the tree |
| inter_len_var | variance of internal branch lengths in the tree |

*Continued on next page*

Table S4 – *Continued from previous page*

| Variable name | Description |
|---|---|
| ter_len_mean | mean of terminal branch lengths in the tree |
| ter_len_var | variance of terminal branch lengths in the tree |
| supp_mean | mean of support values at nodes in the tree |
| rcv | relative composition variability (as described by Phillips and Penny (2003)) |
| treeness_o_rcv | 'treeness' devided by 'rcv' (as described by Phillips and Penny (2003)) |
| saturation | slope of sequence pairs substitution-distance correlation (as described by Philippe et al. (2011)) |
| LB_std | standard deviation of the long branch score per sequence (as described by Struck (2014)) |
| gc_mean_pos1 | mean of GC-content per sequence at codon position 1 |
| gc_var_pos1 | variance of GC-content per sequence at codon position 1 |
| gap_mean_pos1 | mean of gap percentage per sequence at codon position 1 |
| gap_var_pos1 | variance of gap percentage per sequence at codon position 1 |
| gc_mean_pos2 | mean of GC-content per sequence at codon position 2 |
| gc_var_pos2 | variance of GC-content per sequence at codon position 2 |
| gap_mean_pos2 | mean of gap percentage per sequence at codon position 2 |
| gap_var_pos2 | variance of gap percentage per sequence at codon position 2 |
| gc_mean_pos3 | mean of GC-content per sequence at codon position 3 |
| gc_var_pos3 | variance of GC-content per sequence at codon position 3 |
| gap_mean_pos3 | mean of gap percentage per sequence at codon position 3 |
| gap_var_pos3 | variance of gap percentage per sequence at codon position 3 |

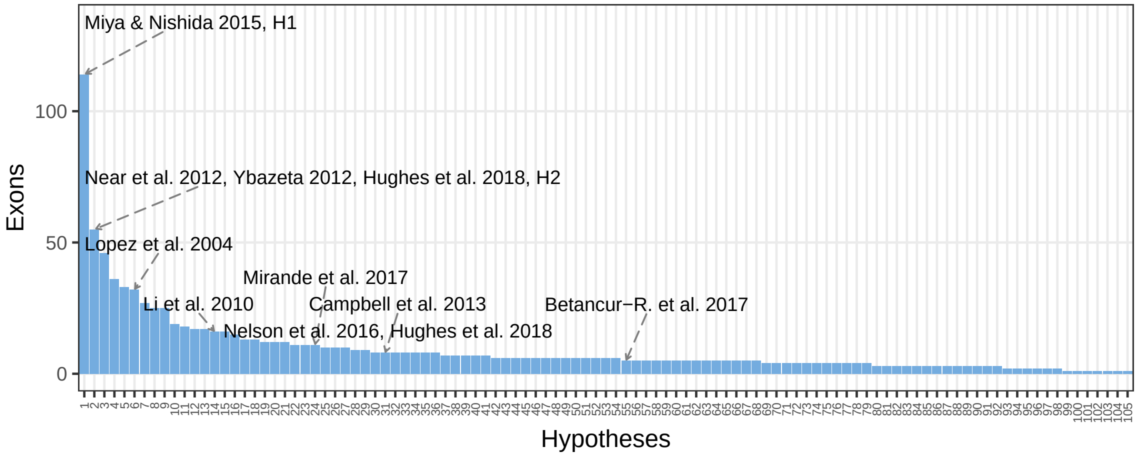

Figure S2: The frequency distribution of 'rank 1' exon loci for each GGI hypothesis in the Protacanthopterygii dataset. Bars represent the frequency of the preferred hypothesis (i.e., the one with the lowest AU p-value) for exon loci based on the AU p-value. Arrows above the bars indicate previous studies that support a given GGI hypothesis.

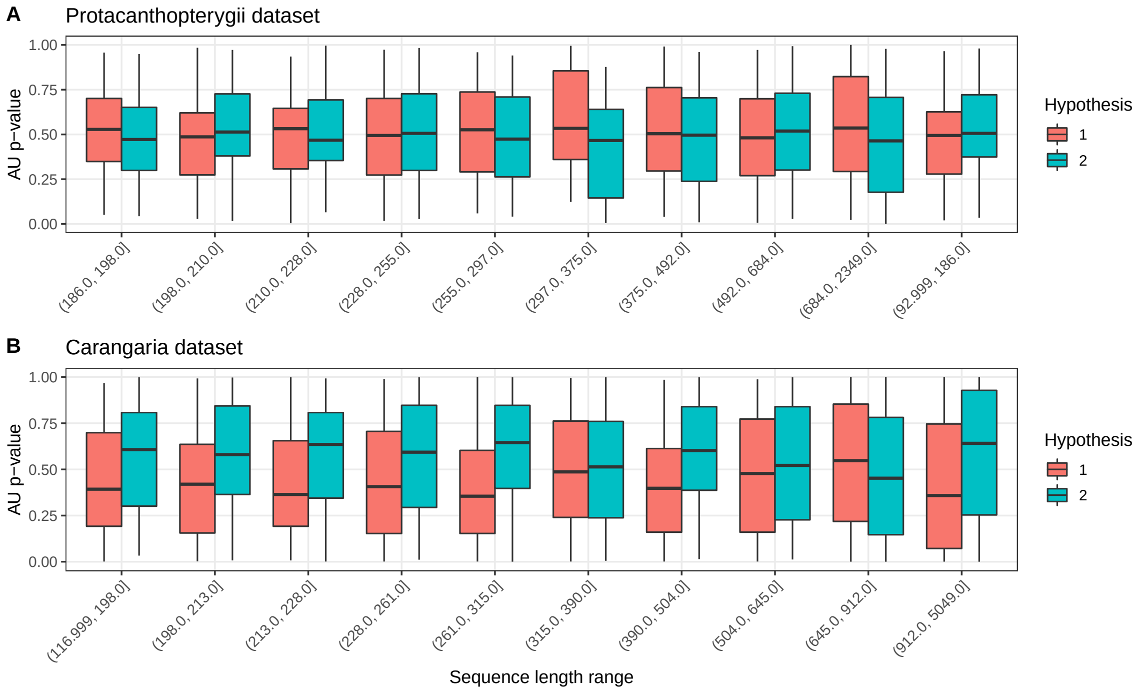

Figure S3: Distribution of both H1 and H2 AU p-values along different range of sequence lengths in both datasets. **A** shows the Protacanthopterygii dataset and **B** shows the Carangaria dataset. For the Protacanthopterygii dataset, the number of loci per bin ranged from 154 to 190. Likewise, in the Carangaria dataset, the count ranged from 164 to 226.

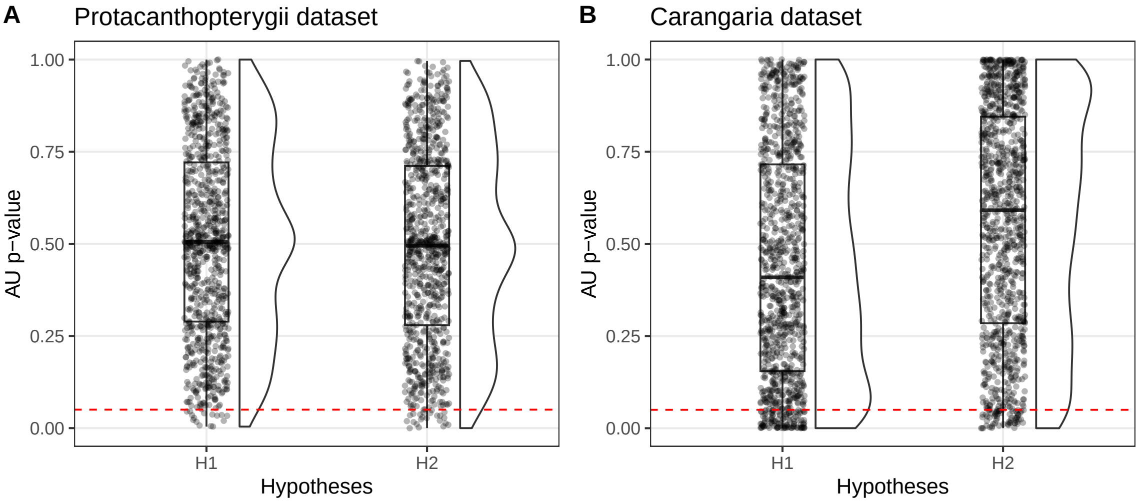

Figure S4: Distribution of both H1 and H2 AU p-values in both datasets. **A** shows the Protacanthopterygii dataset and **B** shows the Carangaria dataset. The dashed line indicates a p-value of 0.05.
